# Reactivation of an errantivirus in *Drosophila* ovarian somatic tissue: from germline invasion to taming

**DOI:** 10.1101/2022.08.29.505639

**Authors:** Marianne Yoth, Stéphanie Maupetit-Méhouas, Abdou Akkouche, Nathalie Gueguen, Benjamin Bertin, Silke Jensen, Emilie Brasset

## Abstract

Most Drosophila transposable elements (TEs) are LTR retrotransposons, some of which belong to the genus Errantivirus and share structural and functional characteristics with vertebrate endogenous retroviruses (ERVs). These virus-derived elements occupy a large part of the genome, but it is unclear whether and how they can be reactivated and if they retain their replication capacity. We created conditions where control of the Drosophila *ZAM* errantivirus through the piRNA pathway was abolished leading to its reactivation in real time in somatic gonadal cells. We show that ZAM may remain active in these cells indicating that errantiviruses may hide from the efficient germline piRNA pathway by being expressed exclusively in somatic cells. After reactivation, *ZAM* invaded the oocytes and severe fertility defects were observed. The germline then set up its own adaptive genomic immune response against the constantly invading errantivirus, restricting invasion and restoring fertility. Our results not only highlight how errantiviruses and their host adapt to each other but also reveal a time window during oogenesis that may be favourable for viral germline invasion and endogenization.

## Introduction

In vertebrates, many LTR retrotransposons belong to the family of endogenous retroviruses (ERVs). ERVs are a class of transposable elements (TEs) that are closely related to infectious retroviruses. Retroviruses are a family of RNA viruses defined by their ability to reverse transcribe their RNA genome into DNA which is then integrated into the host cell genome. While most retroviruses infect somatic cells, occasionally they may infect germ cells and if integrated in the germline genome, they may emerge as ERVs: they become permanent elements in the host genome and are vertically transmitted. ERVs are then capable of replicating themselves and integrating at new genomic loci, which contributes to their success in invading vertebrate genomes. In Human, ERV-related sequences occupy 9% of the genome, in mice 12%, in rat 10.2%, in microbats 5.3%, in marmosets 7.5% (http://repeatmasker.org/genomicDatasets/RMGenomicDatasets.html). Each new ERV copy then evolves independently and can be at the origin of the emergence of a new TE family. In mice, IAPE and IAP ERVs illustrate the evolutionary history of ERVs ^1^. Indeed, IAPEs are still retroviruses capable of infecting other cells, after an extracellular passage, whereas the highly repeated IAPs, derived from IAPEs by losing the envelope gene (env), are strictly intracellular, evolving within a single cell.

It is crucial to note that TEs, can only be maintained in a species if they are able to transpose into the germline genome. If they do not, at least from time to time, they will accumulate mutations until there are no functional copies left and the TE family dies out. Some TEs gained cell and developmental stage specific expression ^2–6^. TEs that have acquired the capacity to be expressed exclusively in somatic cells must therefore maintain their ability to invade germ cells to survive.

ERVs are TEs, and although the presence of TEs in the genome has been shown to provide some evolutionary benefits (reviewed in ^7^), unregulated TE expression and transposition represent a threat to genome integrity and host fitness. Thus, under physiological conditions, most TEs are repressed by epigenetic mechanisms and maintained in a dormant state. In metazoan gonads, TE expression and mobilization is restrained by PIWI-interacting RNAs (piRNAs) (reviewed in ^8, 9^). The piRNA pathway has been extensively studied in the *Drosophila melanogaster* ovary that comprises about 16 ovarioles, each of which contains a succession of follicles composed of germ cells surrounded by somatic follicle cells ^10^ . piRNAs originate from specific source loci, the piRNA clusters ^11^. In Drosophila, the piRNA clusters expressed in the germline are particularly numerous and diverse while there is only one major piRNA cluster expressed in gonadal somatic cells, that is *flamenco* (*flam*) ^12–16^.

Within the Drosophila genome, there is a specific group of TEs that share structural and functional characteristics with vertebrate ERVs. These insect LTR transposons are classified as errantiviruses and belong to the Metaviridae family. In *Drosophila melanogaster* some errantiviruses have been shown to have the capacity to move from cell to cell. These include TEs such as *ZAM, Idefix* or *Gypsy* whose expression is restricted to the somatic follicle cells of the ovary^17–24^. The intercellular invasion capacity may pose a particular threat, especially if they are able to cross the so-called Weismann barrier separating somatic and germ cells (reviewed in^25^). In the germ cells, TE expression and transposition are known to be controlled by the piRNA pathway, but research has primarily concentrated on how piRNAs silence TEs that are expressed in the cells where the corresponding piRNAs are produced. The role of piRNAs in the silencing of a TE that is activated in a cell type different from the one where the silencing mechanism must take place remains unclear. Consequently, whether these TEs can escape the robust epigenetic control mediated by piRNAs when arriving in the germline and how they are tamed by the host over time remained to be puzzled out.

The *ZAM* errantivirus was discovered through its uncontrolled activity in an unstable line, RevI- H2, about 25 years ago ^26, 27^. *ZAM* has 3 intact ORFs encoding proteins whose function is analogous to that of the Gag, Pol and Env proteins of ERVs or retroviruses ^26^. In the RevI-H2 Drosophila line bearing a large deletion in the *flamenco* piRNA cluster, *ZAM* had become active and is specifically expressed in gonadal somatic cells ^14, 17^. Moreover, *ZAM* inserted into multiple new loci in the RevI-H2 line, including a germline-specific piRNA cluster, resulting in unexpected production of germline piRNAs that map to *ZAM* ^26–28^. The biological role of these germline *ZAM*-mapping piRNAs remained unexplored.

Since these studies, *ZAM* has emerged as a key model for studying host-TE interactions and the relationship between somatic cells and the germline during TE invasion.

By analyzing different conditions of *ZAM* reactivation, we aimed here to study what happens in real time when an errantivirus is reactivated *de novo* and how the host responds. Specifically, we sought to elucidate whether piRNAs produced in the germline counteract TE invasion from the somatic cells or whether TE transcripts are protected from degradation by virus-like particles when arriving in the germline.

We show that in the RevI-H2 line, *ZAM* is still expressed in the follicle cells but it doesn’t invade the germline while *ZAM* copies with invasive capacities have been maintained in the genome. To recreate the initial unstable condition, we *de novo* reactivated *ZAM* in the ovarian follicle cells, either by soma-specific knock-down of the piRNA pathway, or by deleting a *ZAM* copy in the somatic *flamenco* piRNA cluster, while keeping the piRNA pathway fully functional. *De novo ZAM* reactivation in the follicle cells led to a massive *ZAM* invasion deep into the adjacent oocyte and its ooplasm. We demonstrate that this invasion could be impeded by the expression of *de novo ZAM*-targeting piRNAs produced in the germ cells themselves demonstrating that these piRNAs are functional against the native *ZAM*. These results show that by being expressed exclusively in somatic cells, errantiviruses may evade the control by the very efficient germinal piRNA pathway and remain active for long periods of time. The challenged germ cells then mount an adaptive genomic immune response to tackle the permanent invasion.

## Results

### In an ancient *flamenco* mutant line, ZAM is still expressed, but the line is nevertheless stable

To investigate the history of *ZAM* transposition dynamics, we monitored *ZAM* copies in the RevI-H2i2 line, an isogenic line that was recently derived from the parental flam mutant RevI- H2 line that has more than 25 years of laboratory history. It had previously been reported that the RevI-H2 line was unstable and that *ZAM* actively transposed in this line ^27^. The *ZAM* instability in RevI-H2 had been linked to a deletion of a large part of the *flam* piRNA cluster. This deletion spanned over more than 120 kb and comprised many different TE relics, including more or less recent copies, and all *ZAM*-related copies of the *flam* locus ^14, 15^.

In the RevI-H2i2 genome, Oxford Nanopore Technology (ONT) genome sequencing revealed 18 *ZAM* copies (Fig. 1a and Table 1 for coordinates). Only one of these *ZAM* copies was also present in the reference genome (http://flybase.org). The other 17 new insertions were very similar to the *ZAM* reference element identified in the initial RevI-H2 line ^26^ and we could clearly localize 14 of the 17 new *ZAM* insertions. Interestingly, ONT long-read sequencing, allowed identifying different *ZAM* variants. Specifically, six were full-length *ZAM* elements (ZAM-fl), one, named *ZAM-v1,* harbored a deleted 5’-UTR (5’ untranslated region), two, named *ZAM-v3*, had a 303 bp deletion within the 5’-UTR, four had large internal deletions in the coding regions, and four were full-length insertions with a deletion at the C-terminal end of the *pol* gene of positions 5494 to 6120 in the *ZAM* internal sequence ^26^ (Repbase *ZAM*_I, https://www.girinst.org/repbase/29). This deleted part of the *pol* gene does not correspond to any known protein domain and the fact that there are five identical copies of this *ZAM* at different locations in the RevI-H2i2 genome indicates that this *ZAM* variant, which we named *ZAM-v2*, is competent for transposition. The full-length *ZAM*, *ZAM-v1* and *ZAM-v3* copies all potentially encode Gag, Pol and Env proteins (Fig. 1a).

**Figure 1:**
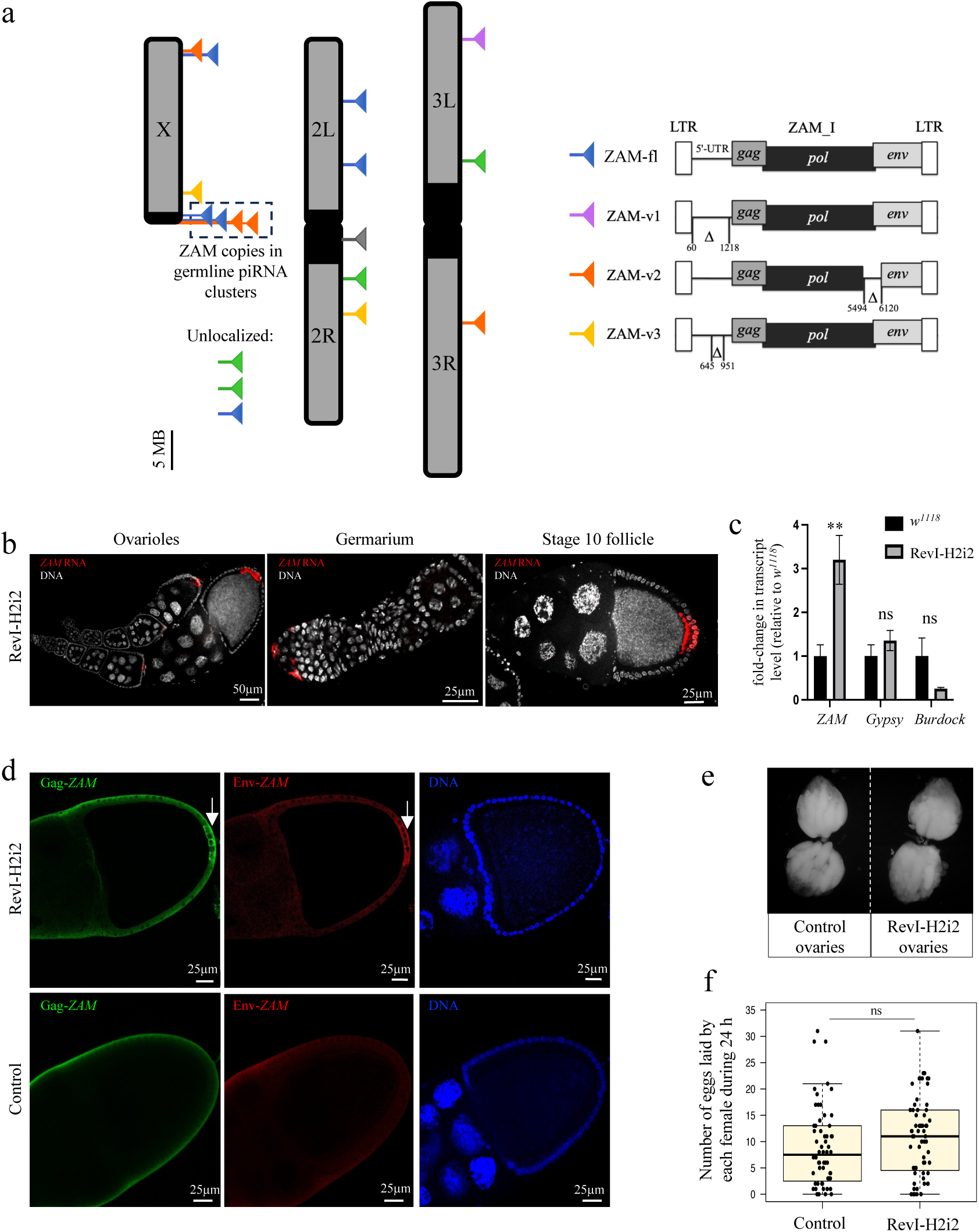
*ZAM* is expressed in follicle cells in the RevI-H2i2 isogenic line but the line is stable. **a**. *ZAM* elements in the RevI-H2i2 genome. Each triangle represents a *ZAM* insertion. The different ZAM-variants are presented in the illustration on the right. The positions refer to *ZAM* internal sequences (*ZAM*_I in RepBase). Blue: full-length *ZAM* elements (ZAM-fl); violet: *ZAM* variants *ZAM-v1* with deleted 5’-UTR; orange: *ZAM*-variants *ZAM-v2*, in which positions 5494-6120 are deleted; yellow: ZAM-variants ZAM-v3 with a 303 bp deletion in the 5’-UTR; green: *ZAM* with internal deletions other than *ZAM* variants above; gray: full-length *ZAM* copy also present in the reference Release 6 genome (http://flybase.org). The dotted box indicates four *ZAM* copies on the X chromosome inserted in referenced germline piRNA clusters (Cluster 9 and 13). No *ZAM* insertion was found in chromosome 4 (1.35 Mb, not shown). For coordinates and details see Supplementary Table 1. **b**. Projection of confocal images of ovarioles, germarium, and stage 10 follicles showing *ZAM* expression by smRNA FISH (in red) in RevI- H2i2 ovaries. DNA was stained with DAPI (white). **c**. Fold-change in the steady-state *ZAM*, *Gypsy* and *Burdock* RNA levels for RevI-H2i2 ovaries compared with *w*^1118^ ovaries (control), quantified by RT-qPCR (primer sequences in Supplementary Table 5). At least three biological replicates and three technical replicates were used. \*\**p-value* <0.01; ns, not-significant (*p-value* >0.05) (Mann-Whitney test). Error bars indicate the standard deviation (SD). **d**. Confocal sections of stage 10 egg chambers of RevI-H2i2 and *w*^1118^ control line showing *ZAM*-encoded Gag (green) and Env (red) proteins. DNA was stained with DAPI (white). **e**. Morphology of control and RevI-H2i2 ovaries. **f.** Box plot displaying the number of eggs laid per fly per day by control and RevI-H2i2 females. Each dot represents an individual female. The midline indicates the median value, and the box shows the first and third quartile. Error bars indicate the SD. ns, not-significant (*p-value* >0.05) (Mann-Whitney test).

In the initial RevI-H2 line, a *ZAM* insertion was found in the germline piRNA cluster *9*, which is close to the X chromosome centromere ^28^. Here, in the RevI-H2i2 line, thanks to long-read sequencing, we could identify *ZAM* insertions in repeated sequences, such as piRNA clusters, more accurately. We now identified three *ZAM* insertions in piRNA cluster *9* (one *ZAM-fl* and two *ZAM-v2*), and also a *ZAM-fl* insertion, which is localized in either piRNA cluster *13* or cluster *56* (uncertainty is due to flanking repeats, piRNA clusters as in ^30^) (Fig. 1a). We reanalyzed the Illumina sequencing data of the initial RevI-H2 line and found evidence that all these *ZAM* insertions in piRNA cluster 9 and 13 or 56 were already present ten years ago, and that at least 14 of the *ZAM* insertions in the RevI-H2i2 line were already present in the parental RevI-H2 line. This result indicates that the genomic *ZAM* profile is the same as ten years ago and suggests that no new *ZAM* insertions occurred since then in the germline.

Strikingly, no new *ZAM* insertion was detected in the *flam* piRNA cluster. Thus, in the RevI- H2i2 line, no *ZAM* piRNA can be produced from *flam*, the major somatic piRNA source locus.

Interestingly, single-molecule RNA fluorescence *in situ* hybridization (smRNA FISH) revealed the presence of *ZAM* RNA in RevI-H2i2 follicle cells, demonstrating that *ZAM* is not silenced in these somatic cells and that transcriptionally active copies of *ZAM* are still present in the genome. *ZAM* transcripts were produced in follicle cells, starting very early during oogenesis in the germarium. Then, after stage 8, *ZAM* transcripts accumulated in a patch of follicle cells located at the posterior side of the follicles. We did not detect any *ZAM* staining in germ cells (nurse cells or oocytes) (Fig. 1b). RNase A treatment led to complete loss of *ZAM* staining in the follicle cells (Supplementary Fig. 1a). We did not detect any *ZAM* RNA in the Iso1A, *w*^1118^ and *w^IR^*^6^ control ovaries (Supplementary Fig. 1b). Using RT-qPCR, we confirmed that *ZAM* was expressed in the RevI-H2i2 ovaries. Conversely, other TEs such as the soma-specific *Gypsy* and the germline-specific *Burdock*, were not upregulated (Fig. 1c). Interestingly, we observed that, besides the full-length ZAM, at least *ZAM* variants ZAM-v2 and ZAM-v3 are expressed in RevI-H2i2 ovaries (Supplementary Fig. 1c,d). *ZAM* is an errantivirus that encodes Gag, Pol and Env proteins. Using immunostaining, we were able to detect *ZAM* Gag and Env proteins that accumulated in the posterior follicle cells in late-stage follicles (> stage 9) (Fig. 1d). All these results showed that *ZAM* transcripts and proteins are still expressed in the RevI-H2i2 line. Although *ZAM* was expressed in follicle cells, the RevI-H2i2 line was fertile. Indeed, the ovary morphology and the number of eggs laid were comparable in RevI-H2i2 and control females (Fig. 1e, f). Altogether, these results indicate that no silencing mechanism had been set up to repress the *ZAM* errantivirus in follicle cells suggesting that *ZAM* expression has no major deleterious effect in the RevI-H2i2 line and that there is no selective pressure to specifically repress *ZAM* in the follicles cells. These findings suggest that RevI-H2i2 is a stable line although *ZAM* is actively expressed in follicle cells.

### Germline *ZAM-*targeting piRNAs constrain *ZAM* invasion from adjacent somatic cells

Transposition of *ZAM* in the initial RevI-H2 line had occurred in the germline, as attested by the transmission of *ZAM* insertions to the offspring, despite the fact that *ZAM* is specifically and exclusively expressed in the somatic follicle cells. This confirms that the *ZAM* errantivirus, when expressed in follicle cells, transits to the germline to integrate into the germ cell genome. However, we showed that *ZAM* insertions are stabilized in the RevI-H2i2 line. The RevI-H2i2 flies produce high amounts of sense and antisense *ZAM*-derived piRNAs with a ping-pong signature, revealed by an over-representation of 10-nucleotide 5’-overlaps between sense and antisense *ZAM*-derived piRNAs (Fig. 2a), while there was no ping-pong signature for ZAM in diverse control lines ^23, 28^. Ping-pong amplification of piRNAs can only occur in the germline in Drosophila (reviewed in ^31^). Thus, these *ZAM* piRNAs observed in RevI-H2i2 are derived from the germline. Importantly, our previous research has demonstrated that the X chromosome of the RevI-H2 line, which contains the *ZAM* copies inserted in germline piRNA clusters, is both necessary and sufficient to produce *ZAM*-regulating piRNAs ^28^. Additionally, in both RevI-H2 ^28^ and RevI-H2i2 background, a *ZAM* sensor transgene is silenced in the germ cells (Supplementary Fig. 2a). We therefore hypothesized that piRNAs produced in germ cells can thwart *ZAM* invasion from somatic cells to the germline and thus protect the germline from new *ZAM* transposition. However, piRNAs are known to silence TEs in the cells where they are produced and TEs, such as *ZAM*, are not expressed directly in germ cells but arrive from surrounding somatic cells. Moreover, *ZAM* RNA may transit in an encapsulated form and the capacity of piRNAs to target encapsulated RNAs remains unknown.

**Figure 2:**
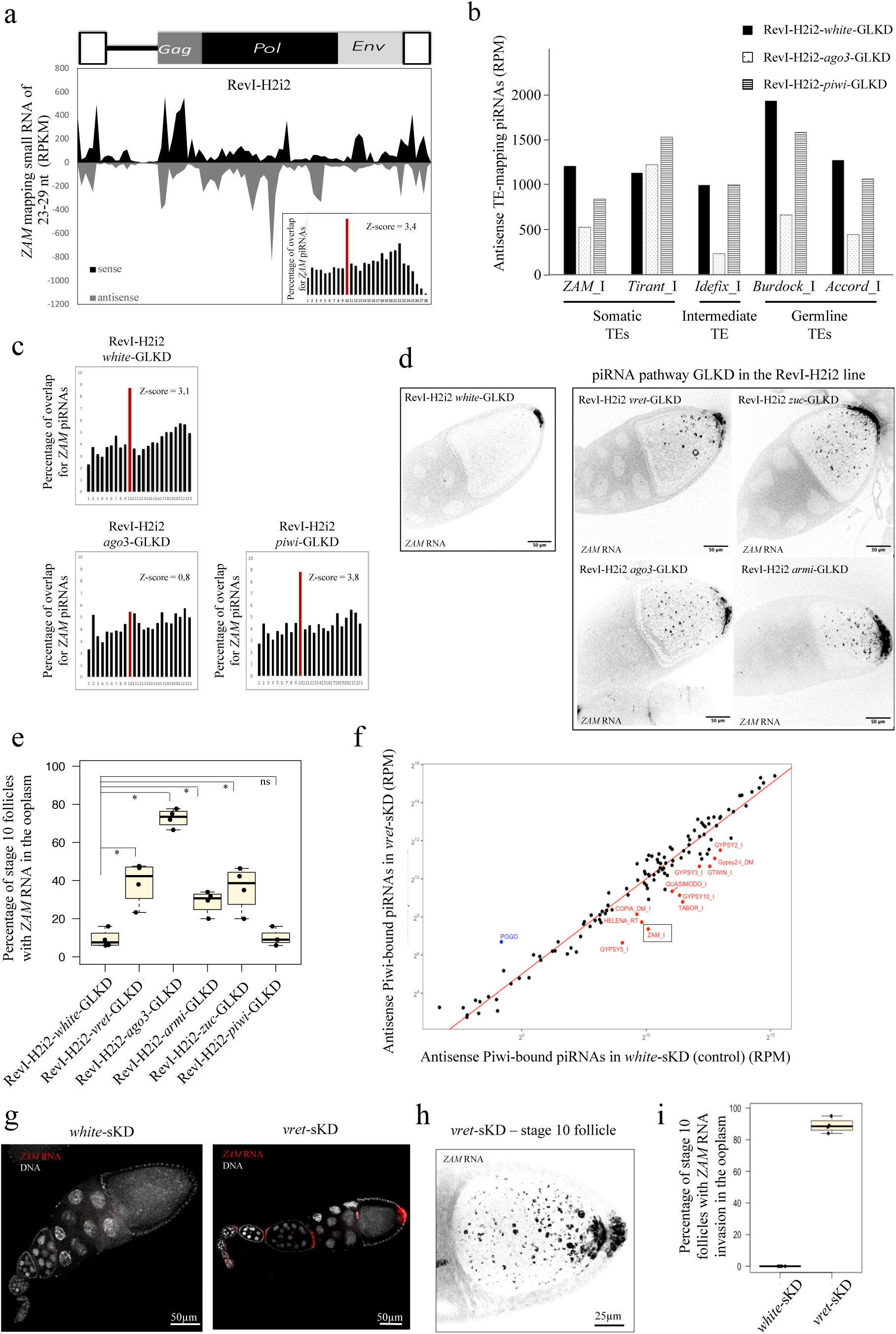
Germline *ZAM* piRNAs produced in RevI-H2i2 germ cells counteract *ZAM* invasion. **a.** Density plot of *ZAM*-mapping regulatory piRNAs along the *ZAM* sequence in RevI-H2i2 ovaries (up to 3 mismatches). In the lower right corner is the histogram showing the percentage of 5’-overlaps between sense and antisense *ZAM*-derived piRNAs (23–29 nt) in RevI-H2i2 ovaries. The proportion of 10nt 5’-overlaps is in red and the corresponding Z-score is indicated. **b.** Antisense piRNAs mapping to TE internal sequences (0-3 mismatches) in the RevI-H2i2 line upon *white-* (control), *ago3-,* or *piwi*-GLKD. Normalized per million of genome-mapping piRNAs. **c.** Histogram showing the percentage of 5’-overlaps between sense and antisense *ZAM*-mapping piRNAs (up to 3 mismatches) in *white-, ago3-,* or *piwi-*GLKD RevI-H2i2 ovaries. The percentage of 10nt overlaps is in red and the corresponding Z-score is indicated. **d.** Color-inverted confocal images of stage 10 egg chambers showing *ZAM* smRNA FISH signal in ovaries of the indicated genotypes. **e.** Bar plot showing the percentage of stage 10 follicles with *ZAM* RNA detected in the ooplasm by smRNA FISH of RevI-H2i2 ovaries with the indicated GLKD. This experiment was done three times and approximately 50 follicles per conditions were observed for each experiment. Error bars indicate the SD. **f.** Scatter plot showing the normalized counts of antisense Piwi-bound piRNAs mapping to individual internal TEs sequences in control ovaries (*white*-sKD) and *vret*-sKD ovaries. Antisense piRNA counts, mapped allowing up to 3 mismatches, were normalized per million of genome-mapping piRNAs (RPM, here in logarithmic scale). TEs in red have a *vret*-sKD/*white*-sKD ratio <0.3. *ZAM* is boxed black. **g.** Confocal images of ovarioles showing *ZAM* smRNA FISH signal (in red) in *white*- (control) and v*ret*-sKD ovaries. DNA was stained with DAPI (white). **h**. Color-inverted confocal projection of stage 10 egg chamber showing *ZAM sm*RNA FISH signal in *vret*-sKD ovaries. **i.** Bar plot showing the percentage of stage 10 follicles with *ZAM* RNA invasion of the ooplasm, assessed by *ZAM* smRNA FISH, of *white*- (control) and *vret*-sKD ovaries. This experiment was done twice and ∼50 follicles per condition were observed for each experiment. Error bars indicate the SD.

To test whether germline ZAM-derived piRNAs were efficient to counteract an invasion coming from surrounding somatic cells, we abolished the piRNA pathway in the germ cells by germline-specific knock-down (GLKD) of piRNA pathway components, in a RevI-H2i2 genetic background producing *ZAM* piRNAs in the germline. It is important to note that, during the genetic crosses performed, only the X chromosome of the RevI-H2i2 was tracked to ensure that the ZAM insertions in the germline piRNA clusters were maintained (crossing schemes presented in Figure S6). We observed that the germline knock-down of proteins of the piRNA pathway, Zucchini (Zuc), Argonaute 3 (Ago3) and Piwi, led to a decrease in the production of *ZAM-*targeting piRNAs, i.e. piRNAs that are complementary and thus antisense to ZAM (Fig. 2b, Supplementary Fig.2b). The decrease in antisense piRNAs was comparable to that observed for germline TEs (i.e., *Burdock* and *Accord*) or intermediate TEs (i.e., *Idefix*), while somatic- TEs such as *Tirant* did not show a similar decrease. Furthermore, in *ago3*-GLKD ovaries, the ping-pong signature for *ZAM* was abolished, as for many other germline-specific TEs, while it was maintained in *piwi-*GLKD ovaries for instance (Fig. 2c, Supplementary Fig. 2c).

We then analyzed the subcellular localization of *ZAM* RNAs by smRNA FISH in the RevI- H2i2 *white*-GLKD control line in which *ZAM*-derived piRNAs are produced in the germline. *ZAM* RNA staining was restricted to follicle cells, only a minor *ZAM* RNA signal was detected in the posterior pole of approximately 10% of stage 10 oocytes (Fig. 2d,e). However, when the piRNA pathway was abolished in RevI-H2i2 germ cells through *vreteno (vret)*, *zuc, ago3* or *armitage (armi) GLKD, ZAM* RNA was no longer restricted to follicle cells, a high signal was also detected in oocytes. *ZAM* RNA spread throughout the ooplasm, but was clearly enriched at the posterior pole of the oocyte, adjacent to the *ZAM*-expressing follicle cells, supporting the fact that *ZAM* RNA originated from follicle cells (Fig. 2d). Actually, 30-70% of stage 10 follicles showed a strong *ZAM* RNA signal in oocytes (Fig. 2e). The strongest invasion phenotype was observed for the RevI-H2i2 *ago3*-GLKD condition where we found a decrease in the production of *ZAM*-derived piRNAs and the loss of the ping-pong signature (Fig. 2b). In this condition, *ZAM* RNA accumulation in the ooplasm was correlated with a strong increase of *ZAM* RNA in total ovaries (RT-qPCR data) (Supplementary Fig. 2d). These results showed that when the piRNA pathway is affected in the germline*, ZAM* RNAs transit from the somatic follicle cells, where they are produced, to the oocyte. Altogether, our data strongly suggest that *ZAM* piRNAs produced by the germline piRNA pathway trigger post-transcriptional silencing of *ZAM* RNAs arriving from follicle cells.

We confirmed this finding using a line where no *ZAM*-derived piRNA is produced in the germline (*w*^1118^ genetic background) and in which we knocked down the piRNA pathway in the follicle cells by somatic knock-down (sKD) of *armi*, *vret*, *piwi* or *yb*. The amount of antisense piRNAs that target soma-specific TEs, including *ZAM,* was strongly decreased (Fig. 2f, Supplementary Fig. 2e, f). RT-qPCR analysis showed that *ZAM* and another soma-specific TE, *Gypsy*, were derepressed, but not the germline-specific *Burdock* (Supplementary Fig. 2g). Importantly, smRNA FISH results revealed that *ZAM* RNAs were not restricted to the posterior follicle cells. Like in ovaries harboring GLKD of the piRNA pathway in RevI-H2i2 background, *ZAM* RNAs were present in oocytes (Fig. 2g,h, Supplementary Fig. 2h). We detected *ZAM* RNAs in the ooplasm in 90% of stage 10 follicles (Fig. 2i). This result confirmed that, when no *ZAM* piRNA is produced in the germline, *ZAM* RNAs expressed in the somatic follicle cells transit to the oocyte.

### *ZAM* RNAs transcribed in follicle cells transit to the oocyte and are conveyed to the embryos in absence of germline ZAM piRNAs

Although *ZAM* is expected to be transcribed only in follicle cells due to its dependence on the Pointed somatic transcription factor ^17, 32^, it was possible that some ZAM copies acquired the capacity to be expressed in the germline over time. Therefore, we wanted to rule out the possibility that *ZAM* RNAs detected in the RevI-H2i2 oocyte originated from germinal nurse cells. If some *ZAM* genomic insertions could be expressed in the germline of the RevI-H2i2 line, then germline depletion of *Piwi*, which is required for the transcriptional gene silencing of TEs, should lead to the transcriptional de-silencing of *ZAM*. We observed that Piwi-GLKD in the RevI-H2i2 ovaries led to the de-silencing of TEs that are capable of transcribing in the germline such as *Burdock*. However, we did not observe any changes in either the *ZAM* RNA level or localization (Fig. 3a,b). Moreover, depletion of *Ago3* induced a substantial accumulation of *ZAM* RNA solely in the ooplasm. In contrast to TEs expressed in the germline (e.g, *Burdock*), we never detected *ZAM* RNA staining in the cytoplasm of the nurse cells at any stage (Fig. 3c). These results strongly support that *ZAM* cannot be expressed in the RevI-H2i2 germ cells and that *ZAM* RNAs detected in oocytes originate from the somatic follicle cells, not nurse cells.

**Figure 3:**
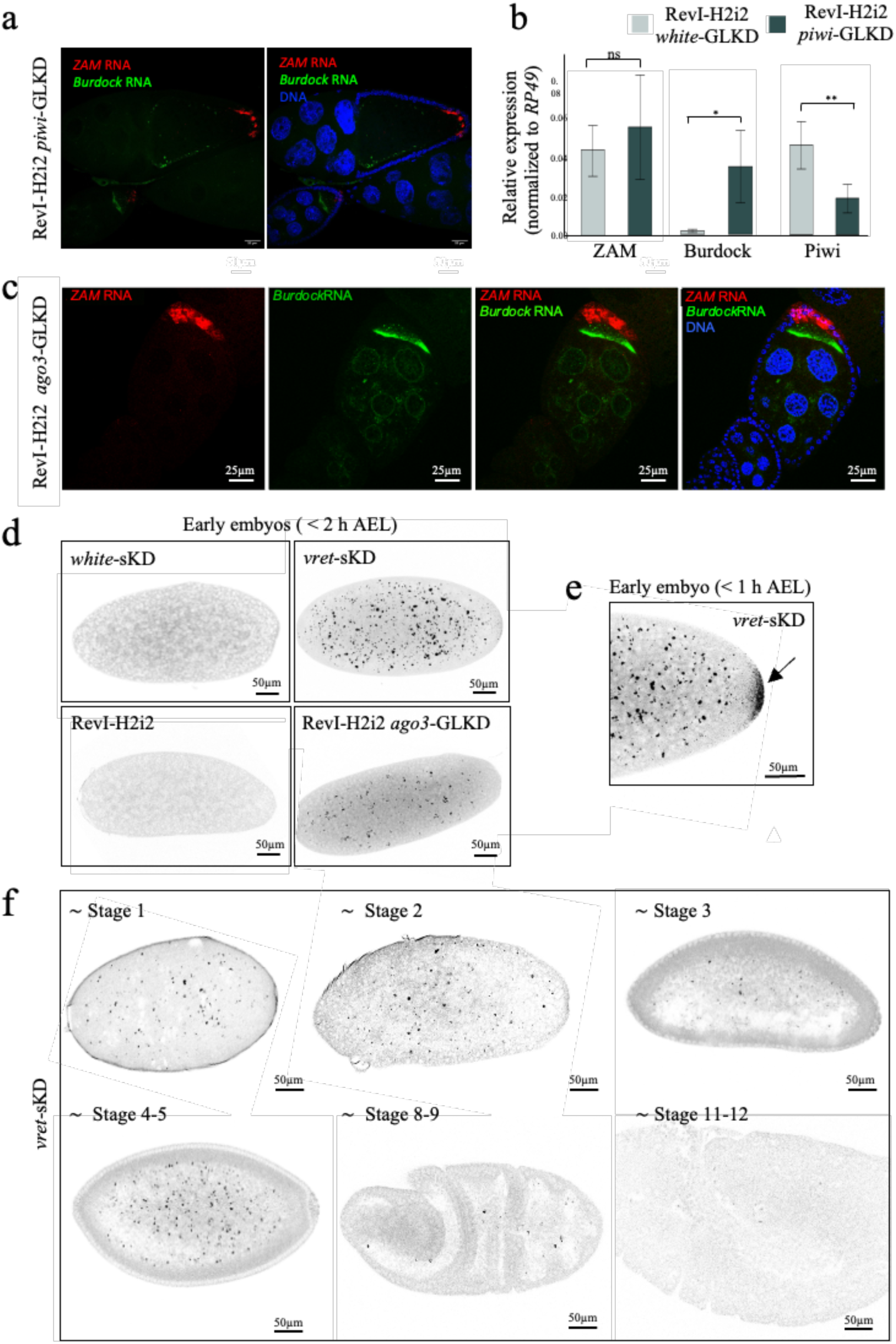
*ZAM* RNAs transcribed in the follicle cells transit to the oocyte and are deposited in early embryos. **a.** Confocal images of stage 10 follicles in the RevI-H2i2 *piwi*-GLKD line. *Burdock* (green) and *ZAM* (red) mRNAs were detected by smRNA FISH. DNA was stained with DAPI (blue). b. Fold-change in the steady-state *ZAM*, *Burdock* and *Piwi* RNA levels for RevI-H2i2 *piwi-* GLKD ovaries compared with RevI-H2i2 *white-*GLKD ovaries (control), quantified by RT-qPCR (primer sequences in Supplementary Table 5). At least three biological replicates and three technical replicates were used. *: *p-*value < 0.05, **: p*-value* <0.01; ns, not significant (*p-value* >0.05) (Mann-Whitney test). Error bars indicate the standard deviation (SD) **c.** Confocal images of ovarioles and of an isolate stage 8 egg chamber in the RevI-H2i2 *ago3*-GLKD line. *Burdock* (green) and *ZAM* (red) mRNAs were detected by smRNA FISH. DNA was stained with DAPI (blue). **d.** Color-inverted confocal images showing *ZAM* smRNA FISH signal in early embryos collected 0-2 hours after egg laying (AEL). Mothers were from the indicated genotype. **e.** Zoom on the posterior part of a *vret*-sKD early embryo collected at 0-1 hours AEL showing *ZAM* RNA accumulation at the posterior pole. **f.** Color-inverted confocal images showing *ZAM* smRNA FISH signal in embryos laid by *vret*-sKD females at the indicated developmental stages.

Throughout the animal kingdom, the first embryonic development stages are controlled by transcripts and proteins deposited by the mother during oogenesis ^33^. In Drosophila, most of the maternal mRNAs are dumped from nurse cells into the oocyte during oogenesis. Although *ZAM* transcripts originate from the somatic follicle cells, when we investigated *ZAM* RNA transmission to the oocyte, we found such transcripts also in early embryos. Indeed, in conditions where *ZAM* RNA had been detected in oocytes, smRNA FISH experiments revealed a strong accumulation of *ZAM* RNAs in early embryos, before the zygotic transition, 0-2 hours after egg laying: from a RevI-H2i2 mother with GLKD of the piRNA pathway, and also from a mother without germinal *ZAM* piRNAs with sKD of the piRNA pathway in follicle cells (Fig. 3d, Supplementary Fig. 3a). Furthermore, *ZAM* RNAs accumulated at the posterior pole of early embryos where future germ cells cellularize (Fig. 3e, Supplementary Fig.3b). We detected *ZAM* RNA in high quantity until stage 5 of embryogenesis and few *ZAM* RNA signal persisting until the cellularization of the blastoderm (∼stage 8-9) (Fig. 3f). In the progeny of the RevI-H2i2 line, we did not detect any *ZAM* RNA, although *ZAM* RNAs were produced in follicle cells of RevI-H2i2 ovaries (Fig. 3d). This confirmed that *ZAM* invasion of the germline does not occur in this line. These data revealed an efficient post-transcriptional silencing of *ZAM* RNAs arriving from follicle cells into the oocyte by piRNAs produced in the germline. This mechanism should limit the transposition of the somatic *ZAM* retrotransposon into the germline genome.

### *De novo ZAM* reactivation leads to massive oocyte invasion and may result in severe fertility defects

Our results demonstrate that an adaptive response can emerge to counteract the invasion of germ cells by an errantivirus. To go deeper, we sought to assess the direct impact of reactivating an errantivirus in real time, before the establishment of any adaptive response by the germline. To conduct this study, we aimed to specifically reactivate *ZAM de novo*. The loss of the piRNA pathway in the somatic follicle cells leads to *ZAM* reactivation, however many other TEs are also desilenced (Supplementary Fig. 2e, g). Moreover, alteration in the expression of piRNA pathway genes also results in severe developmental defects during oogenesis ^34–37^. Therefore, this model cannot be used to specifically analyze the impact of *ZAM* reactivation in ovaries. On the other hand, in the RevI-H2 line where *ZAM* was also reactivated, a large deletion of *flamenco* occurred during non-targeted mutagenesis and an adaptive response had already been set up in the germline. Thus, we chose to create a condition where a single TE is reactivated *de novo* and the piRNA pathway is fully functional.

Using CRISPR-Cas9, we *de novo* deleted the longest *ZAM* copy in the *flam* piRNA cluster in a line carrying the X chromosome of the Iso1A reference line. As in this line, the *flam ZAM* copy is at the genomic position X:21,778,810..21,783,994, we used two guides designed to create a deletion spanning over X:21,777,135..21,784,062 (6926 pb). We named the resulting line *flamΔZAM*. Mapping of genome-unique piRNAs to the *flam* locus highlighted the complete loss of piRNA production in the targeted region compared with the control Iso1A line (Fig. 4a). Conversely, the global production of genome-unique piRNAs mapping upstream and downstream of *ZAM* was not affected. We confirmed that the deletion induced a strong decrease of all piRNAs mapping to the internal regions of the reference *ZAM*. However, piRNAs targeting the *ZAM* LTR were still produced (Fig. 4b). In line with these results, PCR amplification and DNA sequencing of the CRISPR-Cas9 target locus showed that only the internal regions of *ZAM* had been deleted from the *flam* piRNA cluster, resulting in the retention of a solo-LTR at the initial *ZAM* insertion site (Supplementary Fig. 4).

**Figure 4:**
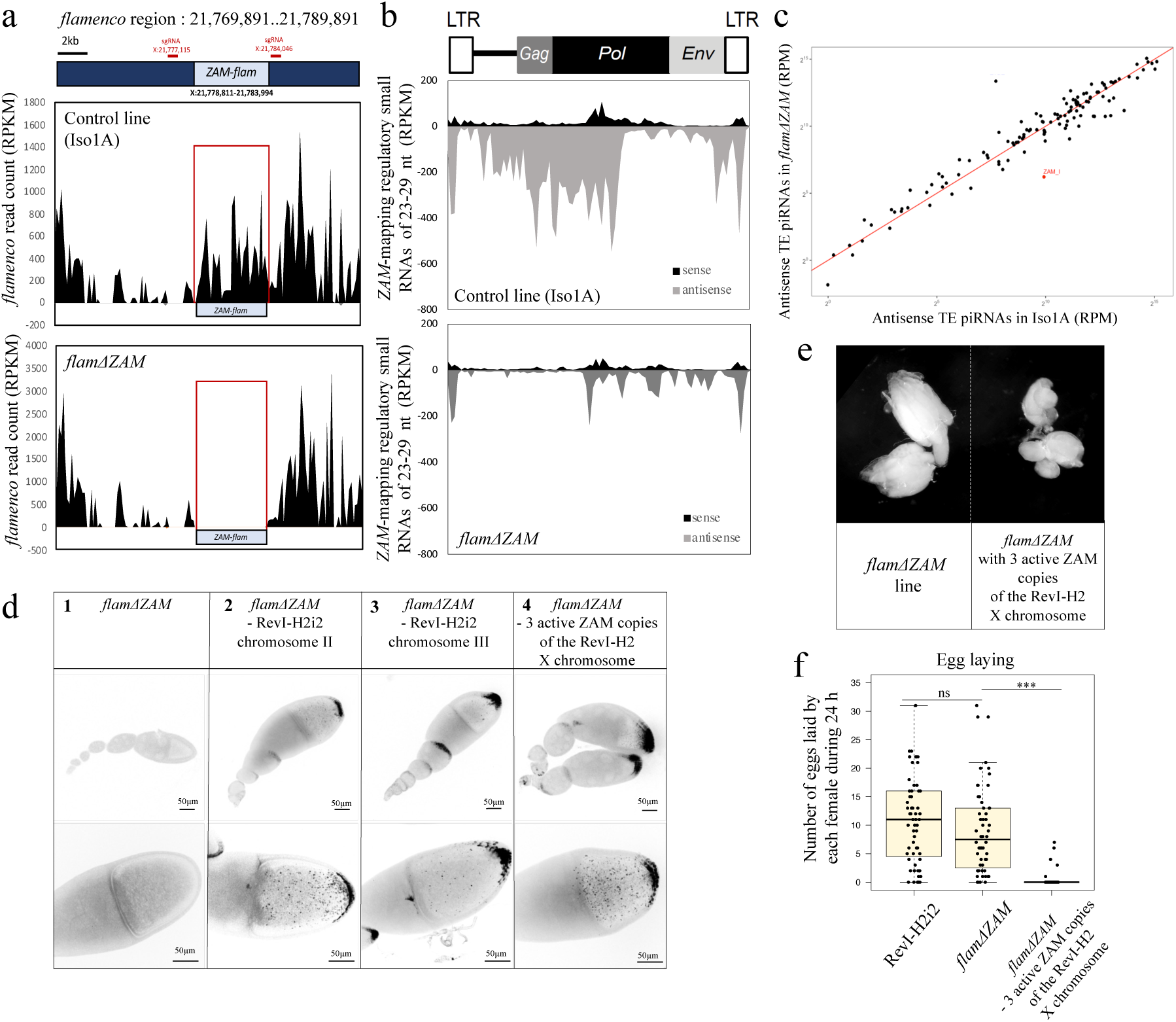
*ZAM* deletion from the *flamenco* piRNA cluster leads to *ZAM* reactivation and oocyte invasion. **a.** Density plots showing genome-unique piRNAs mapping over 20 kb of the *flamenco* piRNA cluster (without mismatch) where the *ZAM* insertion is located (Release 6: X:21,769,891..21,789,891) in the control (Iso1A) and *flam*Δ*ZAM* lines. The position of the *flamenco ZAM* copy *(ZAM-flam)* is indicated. The positions of the sgRNAs used for *ZAM-flam* deletion by CRISPR-Cas9 are shown in red. **b.** Density plot of *ZAM*-mapping regulatory piRNAs along the *ZAM* sequence in control and *flamΔZAM* ovaries (up to 3 mismatches). **c.** Scatter plot showing the normalized counts of antisense regulatory piRNAs mapping to individual internal TE sequences in control ovaries (Iso1A) versus *flamΔZAM* ovaries. Antisense piRNA counts, mapped allowing up to 3 mismatches, were normalized per million of genome-mapping piRNAs (RPM, here in logarithmic scale). TEs in red have a *flamΔZAM*/*Iso1A* ratio <0.3. **d.** Color-inverted confocal images of ovarioles (upper panels) and stage 10 egg chambers (lower panels) from the indicated genotypes showing *ZAM* smRNA FISH signal. **e.** Ovaries of the *flamΔZAM* line and of the *flamΔZAM* line with three recent *ZAM* copies introduced by genetic crossing and X chromosome recombination with the RevI-H2 line. **f.** Box plot displaying the number of eggs laid per fly per day by RevI-H2i2, *flamΔZAM and flamΔZAM* females with three recent *ZAM* copies. Each dot represents an individual female. The midline indicates the median value, and the box shows the first and third quartile. Error bars indicate the SD. \*\*\**p-value* <0.001; ns, not-significant (*p-value* >0.05) (Mann-Whitney test).

In the *flamΔZAM line*, only the production of *ZAM*-derived antisense piRNAs was strongly altered, whereas the production of antisense piRNAs mapping to other TEs was not affected (Fig. 4c). This shows that the *ZAM* deletion from *flam* impairs *ZAM*-derived piRNA production, but does not affect the global piRNA production in ovaries. In sum, in the *flamΔZAM* line, the piRNA pathway was functional and almost no piRNAs that could target the internal *ZAM* sequences were produced.

We then analyzed *ZAM* RNA expression by smRNA FISH in *flamΔZAM* ovaries. Surprisingly, we did not detect any *ZAM* expression (Fig. 4d- panel 1). Based on this observation, we hypothesized that either there was no functional *ZAM* copy in the *flamΔZAM* line, or that the *ZAM* solo-LTR retained in *flam* was sufficient to silence *ZAM* expression. To check this, we exchanged chromosome II or III of the *flamΔZAM* line with chromosome II or III of the RevI- H2i2 line that each contains three or two recently transposed potentially functional *ZAM* copies, respectively. We made sure not to introduce the ZAM copies located in germline piRNA clusters of the X chromosome of the RevI-H2i2 line that are involved in the production of ZAM-regulating piRNAs in the germline. smRNA FISH revealed that *ZAM* RNAs were strongly expressed when these *ZAM* copies were added to the genome of the *flamΔZAM* line (Fig. 4d – panels 2 and 3). We also observed a strong invasion of oocytes with chromosome II, where approximately 70% of stage 10 follicles presented *ZAM* RNAs in the ooplasm, and moderate invasion with chromosome III. Similarly, we observed strong invasion of the oocyte when we added three euchromatic *ZAM* copies located on the telomere side of the X chromosome of the RevI-H2 line to the *flamΔZAM* genome (Fig. 4d – panel 4). Overall, our findings show that in the absence of *ZAM*-derived piRNAs in the somatic follicle cells and in the germline, transcripts from functional *ZAM* copies massively invade the oocytes. Additionally, our results demonstrate that the presence of a solo-LTR of *ZAM* (454 bp) in the *flam* piRNA cluster of the *flamΔZAM* line was not sufficient to silence *ZAM* expression in follicle cells (Supplementary Fig. 4a). The three *ZAM* copies added to the X chromosome of the *flamΔZAM* line consist of one full length *ZAM* insertion (*ZAM-fl*), one *ZAM-v2* and one *ZAM-v3*. Utilizing the sequence specificity of each insertion, we conducted RT-PCR analysis to determine which copies were expressed. The results revealed that *ZAM-fl*, *ZAM-v2* and *ZAM- v3* copies are expressed in RevI-H2i2 ovaries as well as when joined to the *flamΔZAM* genome (Supplementary Fig, 4b).

An important observation was that the *flamΔZAM* flies that contain the three functional *ZAM* copies on the X chromosome displayed atrophied ovaries and reduced fertility compared with the control *flamΔZAM* flies that lack functional *ZAM* copies (Fig. 4e,f). Specifically, 90% of *flamΔZAM* females with active *ZAM* copies did not lay eggs. These data indicate that the reactivation of one single errantivirus in a patch of follicle cells can induce severe fertility defects. These results suggest that when *ZAM* piRNAs are produced in the germline, they contribute to ensure not only genome integrity but also fertility by protecting the oocytes against *ZAM* invasion.

### No initial small RNA response takes place after *ZAM* reactivation

The reactivation of an errantivirus, such as *ZAM,* in a genetic context where the piRNA pathway is functional provides a unique opportunity to examine the genomic immune response to TE reactivation and germline invasion. piRNAs and siRNAs are the two main classes of small RNAs produced to control the expression of transposable elements and counteract viral infection, respectively. Moreover, invading viral RNAs can be processed to small RNAs to initiate a primary response ^38, 39^. To characterize the small RNA response upon *ZAM* reactivation, we sequenced small RNAs that are complexed with Argonaute proteins (named here “regulatory” piRNAs) from *flamΔZAM* ovaries containing two or four potentially functional *ZAM* copies. In the *flamΔZAM* line, there were no *de novo ZAM* piRNAs (23 to 29 nt) produced upon *ZAM* reactivation, despite germline invasion. Indeed, only few sense and antisense *ZAM* piRNAs were detected in the ovaries of the *flamΔZAM* line that contained functional *ZAM* copies (Fig. 5a, Supplementary Fig. 5a). Moreover, this lack of piRNA production concerned only *ZAM*. The level of regulatory piRNAs targeting other TEs was not reduced (Fig. 5b, Supplementary Fig. 5b).

**Figure 5:**
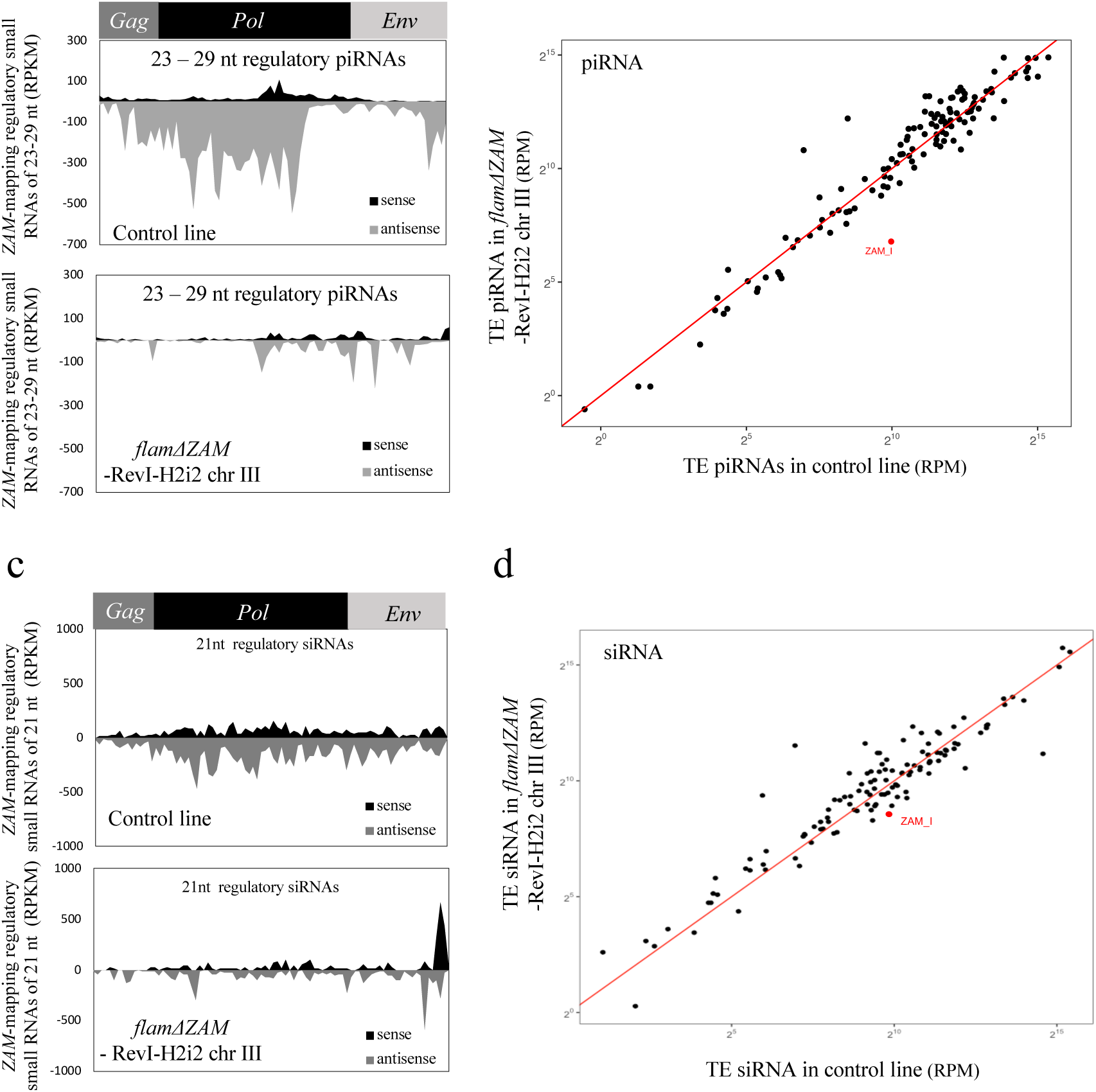
No initial piRNA and siRNA response after *ZAM* retrotransposon reactivation. **a.** Density plot of regulatory piRNAs along the *ZAM* internal sequence in ovaries from a control line (*white*-sKD) and from the *flamΔZAM*-RevI-H2i2 chr III line (up to 3 mismatches). **b.** Scatter plots showing the normalized counts of regulatory piRNAs mapping to individual internal TE sequences in control ovaries (*white-sKD*) versus *flamΔZAM*-RevI-H2i2 chr III ovaries. piRNA counts, mapped allowing up to 3 mismatches, were normalized per million of genome-mapping piRNAs (RPM, here in logarithmic scale). **c**. Density plot of *ZAM*-mapping regulatory siRNAs (21nt small-RNAs complexed with Argonaute proteins) along the *ZAM* internal sequence produced in control (*white*-sKD) and in *flamΔZAM* RevI-H2i2 chr III ovaries (0-1 mismatch). **d**. Scatter plot showing the normalized counts of antisense regulatory siRNAs mapping to individual internal TE sequences in control ovaries (*white-sKD*) versus *flam*Δ*ZAM* RevI-H2i2 chr III ovaries. Antisense siRNA counts, mapped allowing up to 1 mismatch, were normalized per million of genome-mapping siRNAs (RPM, here in logarithmic scale).

We sequenced total small RNAs in *vret-* or *yb*-sKD ovaries where *ZAM* was also reactivated. Surprisingly we observed sense *ZAM*-mapping small RNAs of various sizes, including 23-29 nt small RNAs that are not produced in a *white*-sKD control condition (Supplementary Fig. 5c). However, the sense 23-29 nt *ZAM*-derived small RNAs detected in *yb*- and *vret*-sKD ovaries did not display uridine bias at the 5’ end, a feature of mature primary piRNAs (Supplementary Fig. 5d). Furthermore, these sense *ZAM*-mapping small RNAs were not immunoprecipitated with Piwi proteins. We detected almost no Piwi-bound sense or antisense *ZAM*-mapping piRNAs in *vret-* or *yb-*sKD ovaries (Supplementary Fig. 5e). In addition, no sense regulatory *ZAM* piRNAs complexed with Argonaute proteins were detected in *vret-*sKD ovaries (Supplementary Fig. 2f). These results suggest that the sense *ZAM*-mapping small RNAs are most likely degradation products of excess *ZAM* sense transcripts and not piRNAs. Thus, in these different conditions, we did not detect any primary piRNA response either in the somatic cells where *ZAM* is expressed or in the oocyte that it invades.

Given that the *ZAM* errantivirus shares some structural and functional characteristics with retroviruses, we also asked whether *ZAM* reactivation could trigger a siRNA response. For this, we analyzed the production of 21 nt small RNAs complexed with Argonaute proteins (“regulatory” siRNAs). Similar levels of *ZAM* 21 nt small RNAs were produced in a control line and in the *flamΔZAM* line with functional *ZAM* copies (Fig. 5c). Globally, the production of 21 nt small RNAs in this line was comparable to the control for all TEs, including *ZAM* (Fig. 5d). *ZAM*-mapping 21 nt small RNAs were slightly increased only in the *vret*-sKD condition, compared to the *white*-sKD control line without *ZAM* expression (Supplementary Fig. 5f). Overall, our results suggest that no or very few siRNAs that could protect ovaries from *ZAM* activity are produced upon activation of this errantivirus in follicle cells.

Therefore, we propose that no innate small RNA response is triggered in the lines where *ZAM* is reactivated, suggesting that *ZAM* will not be brought under control until the establishment of specific adaptive immunity and immune memory.

## Discussion

The discovery that piRNAs produced in germ cells not only restrict TE expression in the germline itself but also counteract the invasion of errantiviruses from adjacent cells reveals a novel role of piRNAs produced locally as an effective defense mechanism at the tissue scale. When piRNAs against *ZAM* are produced in the germline, *ZAM* transcripts are limited to a patch of follicle cells and flies are fertile. Conversely, in the absence of *ZAM*-derived piRNAs in the germline, transcripts from functional *ZAM* copies massively invade the oocytes and are even transmitted to the embryos. At the molecular level, our study revealed that piRNAs produced in nurse cells and dumped into the oocyte can target RNAs produced by follicle cells and delivered to the oocyte. It is tempting to speculate that the *ZAM* RNAs arriving from somatic cells are targeted by complementary antisense piRNAs and degraded through the ping-pong cycle (reviewed in ^31^). This would result in the production of *ZAM* sense piRNAs, thus participating in the amplification of the pool of piRNAs against *ZAM* and strengthening the defense mechanism against the invasion.

Transposition in the germ cell genome is crucial for TE propagation in a population because it allows the vertical transmission of new insertions. At the time of invasion (e.g. in the *flamΔZAM* line), no *ZAM*-derived piRNA was produced in somatic or germ cells. Therefore, this condition could be compared to what happens when a TE first enters a new species by horizontal transfer. In this case, according to several studies, an initial transposition burst occurs that leads to TE accumulation in the genome before the induction of an adaptive response by the host to control transposition ^40–45^. We found at least 17 new *ZAM* insertions in the RevI-H2i2 genome, testifying that *ZAM* actively transposed. Surprisingly, we identified three *ZAM* insertions in the same germline piRNA cluster on the X chromosome. It has been proposed that two different mechanisms can explain the *de novo* piRNA production required to specifically silence a novel invading TE. Indeed, piRNAs can be produced from a new TE insertion into a pre-existing piRNA cluster, and also by stand-alone TE insertions converted into piRNA-producing loci ^46–49^. Interestingly, when active *ZAM* copies are added into a *flamΔZAM* genetic background, these copies do not produce regulatory *ZAM*-derived piRNAs and are not converted into piRNA producing regions. Actually, *ZAM* should be transcriptionally inactive in the germline because *ZAM* expression is regulated by the somatic transcription factor Pointed ^32^. Therefore, although the contribution of individual copies needs to be investigated, our data strongly suggest that germline *ZAM* piRNAs originate at least in part from piRNA clusters. In line with our findings, computer simulations of the TE invasion dynamics suggested that several insertions in piRNA clusters are likely to be required to stop the invasion in the germline ^50^ . Conversely, we found that a single insertion in a somatic piRNA cluster, such as *flamenco,* is sufficient for TE silencing in the somatic follicle cells, as previously suggested ^14^. Indeed, we show here that in follicle cells, the *flamenco* piRNA cluster acts alone as the principal regulator of transposon activity. A precise deletion of the internal part of the *ZAM* TE inserted in *flamenco* leads to complete derepression of functional *ZAM* copies. Surprisingly, it seems that reactivation of this errantivirus in only a patch of somatic cells can lead to severe fertility defects. However, in the germline, the deletion of the three most highly expressed germline piRNA clusters (42AB, 38C, and 20A) affected neither fertility nor TE silencing ^48^. This could be explained by the fact that in the germline, but not in somatic cells, redundancy in piRNA production among different piRNA clusters is observed, with probably multiple piRNA clusters involved in the silencing of the same TE. Moreover, many TE copies have acquired mutations and are non-functional for transcription or for transposition. In the *flamΔZAM* line, although no *ZAM* piRNA was produced, we did not detect any *ZAM* expression. However, when we introduced functional copies of *ZAM*, *ZAM* was expressed demonstrating that depending on the genetic background, the transposon landscape varies and functional copies can be absent or present. In general, the TE content varies considerably among populations ^47, 51^. Interestingly, we identified *ZAM* copies with a deletion that affects the *pol* gene. The copy numbers of this *ZAM* variant *ZAM-v2* and of *ZAM* full-length elements (ZAM-fl) that have transposed in the RevI-H2 line are similar (4 and 6 copies respectively), suggesting that even this *ZAM*-variant can transpose very efficiently. Studies in different organisms have shown that TEs rapidly diversify. For instance, the *mariner* DNA transposon is present in 68 different versions in grass genomes ^52^. Such rapid diversification certainly promotes speciation of new families of active TEs.

One interesting question is how *ZAM* RNAs transit into the oocyte and at which stage during Drosophila oogenesis *ZAM* transposes in germ cells. We noticed that *ZAM* RNAs, although transcribed in follicle cells, were deposited and accumulated at the posterior pole of early embryos, where future germ cells cellularize. *ZAM* might have evolved to preferentially mobilize in the dividing primordial germ cells of the offspring instead of the developing oocyte, which is not in a phase favorable to transposition because of its prolonged arrest in prophase I associated with highly compacted chromatin^53^.

Previous experiments demonstrated that *ZAM* RNAs accumulate in follicle cells when vitellogenesis is defective, suggesting that *ZAM* transmission to the oocytes requires functional vitellogenin trafficking ^20^ (Supplementary Video 1). Moreover, errantiviruses, such as *ZAM* and *Gypsy*, can form pseudo-viral particles in follicle cells when they are expressed ^17, 19, 20, 24^. *ZAM* Gag and Env proteins are expressed in RevI-H2i2 follicle cells. Retroviral envelope glycoproteins undergo proteolytic processing by cellular serine endoproteases (furin and proprotein convertases), in order to produce the two functional subunits, a glycosylated hydrophilic polypeptide (SU) and a transmembrane domain (TM). This step is necessary to achieve protein competence to promote membrane fusion and virus infectivity ^54, 55^. The errantiviral *env* gene, acquired from insect baculovirus ^56^, encodes a protein whose function is analogous to that of retroviral Env protein, which is responsible for infectious properties. Indeed, *ZAM* Env protein is composed of an SU, a TM and an Arg–X–Lys–Arg conserved domain, which is considered to be a furin consensus proteolytic cleavage site ^26, 55, 57^. A recent study demonstrated that intracellular retrotransposition of the *Gypsy* errantivirus can occur in the absence of the Env protein, while intercellular transmission requires a functional Env ^19^. Currently, whether *ZAM* can transpose intracellularly, in follicle cells, and the exact mechanisms (Env protein dependent or independent) used for *ZAM* transmission to the oocyte are unknown.

Once a virus enters the target cells, several scenarios for retrovirus uncoating have been proposed. Recent advances on the HIV-1 strongly suggest an uncoating at the nuclear pore and within the nuclear compartment ^58^. Errantiviruses also encode Gag proteins which are capable of mediating the assembly of virus-like particles. However, there are significant differences between these proteins and retroviral Gag proteins, which might lead to a different uncoating process for errantiviruses ^59^. In our study, we observed the loss of *ZAM* RNA in the oocyte when germline *ZAM*-derived piRNAs are produced. We can hypothesize that the *ZAM* capsid is dissociated soon after entry into the oocyte and that piRNAs can therefore easily target *ZAM* RNAs. It is also possible that *ZAM* RNAs are still encapsidated in the oocyte, but the capsid does not protect the *ZAM* RNA from piRNAs and their associated endonuclease. Previous studies on virus-like particles of the yeast *Ty* retrotransposon suggest that these particles form open structures that leave the RNA at the interior of the capsid accessible to RNase ^60, 61^.

It has been proposed that in different species, an “innate” response at the piRNA or siRNA level can be initiated in the gonads upon retrovirus invasion (^39, 62–67)^. We found that no clear innate small RNA response is triggered upon *ZAM* reactivation in Drosophila ovaries. Total small-RNA sequencing after *ZAM* reactivation revealed the presence of sense *ZAM*-mapping small RNAs, but they lack characteristics of siRNAs or piRNAs. We hypothesize that these sense small RNAs are degradation products generated when there is an excessive amount of *ZAM* RNAs. Altogether, these results show that no initial small RNA response is mounted upon *ZAM* reactivation, leaving follicle cells, where *ZAM* is expressed, and also oocytes unprotected against *ZAM* transposition. Our findings suggest that the piRNA response is a robust response that, in the case of *ZAM*, must be established either in the follicle cells or in the germ cells to efficiently control the invading TE. However, the time required for developing a robust piRNA response to sustainably control TE transposition remains unknown. Lastly, it is thought that the piRNA pathway has no antiviral role in Drosophila ^68^. However, it would be interesting to determine whether this is also true if parts of the viral genome are integrated into a germline piRNA cluster during infection.

Germline and somatic piRNA pathways and their respective piRNA clusters are distinct. There is no ping-pong amplification of piRNAs in the somatic follicle cells and piRNA production in these cells is almost exclusively dependent on a single piRNA cluster, *flamenco*, making it more vulnerable to mutation. A key outcome of our results is that they suggest that not only does the host adapt to the TE but the TE also seems to have adapted to the host: the exclusive expression of *ZAM* in somatic cells allows the TE to “hide” in the follicle cells and invade an unprotected germline after a simple deletion of a short *flam* sequence. The errantivirus *ZAM* thus seems to have adapted to the host by gaining tissue specificity and by adopting an expression niche.

## Methods

### Fly stocks, transgenic lines, and crosses

Flies were maintained at 25°C under a light/dark cycle and 60% humidity. Between 3 and 6 days after hatching, flies were used for experiments. The RNAi lines against components of the piRNA pathway used in this study are listed in **Supplementary Table S2**. Isogenic lines from the RevI-H2 stock were generated by crossing a single female with a single male for five generations. Germline and somatic knock-down have been performed using the nanos-gal4 and the traffic-jam-gal4 driver lines respectively (**Supplementary Table S2)**. The Drosophila line carrying the *ZAM* sensor is described in ^28^. The sensor expresses the GFP reporter gene under the control of an inducible Upstream Activation Sequence promoter (UASp) and harbors a *ZAM* fragment in its 3′-UTR (pGFP-*ZAM*).

### Single-molecule RNA fluorescence *in situ* hybridization (smRNA FISH) in ovaries and embryos

*ZAM* smRNA FISH was performed using 48 probes that target *ZAM* transcripts in a region that is absent in the *ZAM* inserted in the *flamenco* piRNA cluster to detect only transcripts of active *ZAM* copies (sequences in Supplementary Table S3). Ovaries from 3 to 6-day-old flies were dissected in Schneider’s Drosophila Medium and fixed in Fixing Buffer (4% formaldehyde, 0.3% Triton X-100, 1x PBS) for 20 min at room temperature, rinsed three times in 0.3% Triton X-100, once in PBS, and permeabilized in 70% ethanol at 4°C overnight. Permeabilized ovaries were rehydrated in smRNA FISH wash buffer (10% formamide in 2x SSC) for 10 min. Ovaries were resuspended in 50µL hybridization buffer (10% dextran sulfate, 10% formamide in 2x SSC) supplemented with 1µL of smRNA FISH probes. Hybridization was performed with rotation at 37°C overnight. Ovaries were then washed twice with smRNA FISH wash buffer at 37°C for 30 min and twice with 2xSSC solution. Then, DNA was stained with DAPI (1/500 dilution in 2x SSC) at room temperature for 20 min. Ovaries were mounted in 30 µL Vectashield mounting medium and imaged on a Zeiss LSM-980 or Zeiss LSM-800 confocal microscope. The resulting images were processed using FIJI/ImageJ. The RNA signal specificity was confirmed by adding 1 mg/mL of RNase A in 2x SSC for 2h before the hybridization step.

For embryo staining, flies were caged and fed yeast paste. Embryos (0-2h) were collected, dechorionated in 50% bleach for 4 min and rinsed in water. Eggs were fixed in 4% paraformaldehyde/heptane for 20 min, devitellinized by vigorous shaking in 100% methanol and stored in methanol at -20°C. Embryos were rehydrated with 1:1 methanol in 1xPBS, 0.1% Tween-20 for 5 min and twice in 1xPBS, 0.1% Tween-20. Embryos were resuspended in smRNA FISH wash buffer (10% formamide in 2x SSC) for 10 min and then processed for smRNA FISH as described for ovaries. Immunostaining combined with smRNA FISH was performed by adding primary antibodies to the smRNA FISH probes in the hybridization buffer and incubating at 37°C overnight. Embryos were washed twice with smRNA FISH wash buffer and incubated with the secondary antibody for 90 min. After two washes in 2x SSC, embryos were mounted in 30 µL Vectashield mounting medium.

### Immunofluorescence analysis of ovaries and embryos

Ovaries from 3-6-day-old flies were dissected in supplemented Schneider’s medium, ovarioles were separated, and the muscle sheath was removed before fixation to obtain undistorted follicles. Then, ovaries were fixed in 4% formaldehyde/1x PBS/2% Tween-20 for 15 min, rinsed three times with 1x PBS/2% Tween-20, and incubated 2 hours in PBT (1x PBS, 0.1% Triton X-100, 1% BSA). Ovaries were incubated with primary antibodies in PBT at 4°C overnight (antibodies are described in Supplementary Table S4). After three washes in PBT, ovaries were incubated with the corresponding secondary antibodies coupled to Alexa-488 or Cy3 for 90 min. After two washes in 1x PBS, DNA was stained with DAPI (1/500 dilution in 1x PBS) for 20 min. Ovaries were mounted in 30 µL Vectashield mounting medium and imaged on a Zeiss LSM-980 or Zeiss LSM-800 confocal microscope. The resulting images were processed using FIJI/ImageJ.

### RT-qPCR analysis of transposon expression

10-20 pairs of dissected ovaries were homogenized in TRIzol reagent (Ambion). Following DNase I treatment, cDNA was prepared from 1 µg RNA by random priming of total RNA using Superscript IV Reverse Transcriptase (ThermoFisher Scientific). Quantitative PCR was performed with Roche FastStart SYBR Green Master on a the Lightcycler® 480 Instrument. RT-qPCR was used for quantification of transposon mRNA levels (primer sequences in Supplementary Table S5). All experiments were done with three biological replicates and with technical triplicates. Steady-state RNA levels were calculated from the threshold cycle for amplification with the 2^−ΔΔCT^ method; *rp49* was used for normalization.

### Immunoprecipitation of Piwi-ribonucleoprotein complexes

For each genotype, 50 pairs of ovaries from 3-6-day-old flies were dissected, lysed in 1 ml lysis buffer (20 mM HEPES-NaOH at pH 7.0, 150 mM NaCl, 2.5 mM MgCl2, 250 mM sucrose, 0.05% NP40, 0.5% Triton X-100, 1x Roche-Complete). Samples were cleared by centrifugation at 10000 × g at 4°C for 10 min. Extracts were incubated with rotation with rabbit polyclonal anti-Piwi antibodies (4 µg per sample) at 4°C for 4h followed by overnight incubation with Dynabeads^TM^ Protein A (50 µl, Invitrogen, 10002D) at 4°C with rotation. Before incubation, beads were equilibrated in NT2 buffer (50 mM Tris-HCl, pH 7.4, 150 mM NaCl, 1 mM MgCl2, 0.05% NP40). Beads were washed twice with ice-cold NT2 and twice with NT2 in which NaCl concentration was adjusted to 300 mM. Nucleic acids that co-immunoprecipitated with Piwi were isolated by treating beads with 0.7 mg/ml proteinase K in 0.3ml proteinase K buffer (0.5% SDS, 10mM Tris-HCl pH 7.4, 50mM NaCl, 5mM EDTA), followed by phenol/chloroform extraction (phenol at neutral pH) and ethanol precipitation.

### DNA Isolation, Oxford Nanopore Technology sequencing and genome analysis

DNA was extracted from 200 RevI-H2i2 females using the Qiagen Blood & Cell Culture DNA Midi kit. The genomic DNA quality and quantity were evaluated using a Femto Pulse (Agilent) and a Qbit 3.0 (Invitrogen) respectively. Oxford Nanopore Technology (ONT) sequencing was performed by I2BC (Gif-sur-Yvette, France) using five micrograms of genomic DNA. Adapter- trimmed ONT reads were analyzed using NCBI Blastn (https://blast.ncbi.nlm.nih.gov/Blast.cgi) or command-line Blastn (Blastn 2.8.1). The RevI- H2i2 ONT reads containing *ZAM* insertions were detected by Blastn with the reference *ZAM* ^22^, Repbase *ZAM*_I and *ZAM*_LTR sequences (https://www.girinst.org/repbase/, ^29^). The *ZAM* containing reads were then recovered with bedtools getfasta (https://bedtools.readthedocs.io/en/latest/content/tools/getfasta.html, ^69^). The *ZAM* insertion sites where determined using Blastn of the *ZAM* containing reads to the *D. melanogaster* Release 6 genome (http://flybase.org). The absence of empty sites, without a given *ZAM* insertion, was verified by Blastn of the respective empty insertion sites, recovered from the *D. melanogaster* Release 6 genome, to all ONT reads. Sequencing data are available in NCBI GEO database (GSE213456).

#### RNA-sequencing and analysis

3 independent total RNA extractions from 30 ovaries from 3-6-day-old RevI-H2i2 flies using Trizol (Invitrogen) were performed. Strand-specific libraries for RNA sequencing (RNA-Seq) were constructed at BGI. 1 µg of total RNA was used to prepare mRNA library using MGIEasy RNA Library Prep kit and sequenced as PE150 reads on the DNBSEQ G400 sequencer, and adapter-clipped reads were provided by BGI. The data were analyzed on the local Galaxy platform using HISAT2 alignment (Galaxy Version 2.1.0). The reads were first mapped to *ZAM-fl* and to *ZAM-v2* using the “Disable spliced alignment” option. The mapped reads were then aligned to ZAM_I sequence (Repbase) with the default options, i.e. allowing split alignment. Reads were visualized using “Integrative Genomics Viewer” version 2.16.1. The coverage of RNA-seq reads corresponding to *ZAM-v2* was determined as the number of split reads specific to the *ZAM-v2* deletion at positions ZAM_I:5494-6120, normalized to the average coverage of the pointed transcript. Coverage of RNA-seq reads corresponding to ZAM- fl and ZAM-v2 was determined as the number of non-split reads at position 5495, the left edge of the deletion, normalized to the average coverage of the pointed transcript. Only the left edge was used for this purpose, as the right edge of the deletion is also mapped by reads corresponding to ZAM-variants with other internal deletions (green in Figure 1a).

### Small RNA sequencing

Regulatory small RNA extraction was performed as described in ^70^. Briefly, 50 pairs of ovaries from 3-6-day-old flies were lysed and Argonaute-small RNA complexes were isolated using TraPR ion exchange spin columns (Lexogen, Catalog Nr.128.08). Total RNA also was extracted for some samples (80-100 ovaries for each sample) using the classical TRIzol extraction, followed by 2S rRNA depletion and size selection before sequencing. Sequencing was performed by Fasteris SA (Geneva, Switzerland) on an Illumina NextSeq550 instrument (13-15 million reads per sample).

Illumina small RNA sequencing reads were loaded on the small RNA analysis pipeline sRNAPipe ^71^ for mapping to the various genomic sequence categories of the *D. melanogaster* genome (Release 6). For the analysis, 23-29nt genome mappers were selected as piRNAs, and 21nt genome mappers were selected as siRNAs. “Genome-unique” piRNAs were defined as 23-29 reads that mapped uniquely across the reference genome. piRNAs were mapped to TEs allowing up to 3 mismatches, siRNAs allowing up to 1 mismatch, and genome-unique piRNAs to piRNA clusters as defined in ^11^ allowing no mismatch. The window size was of 91nt for the entire *ZAM*, 86 nt for ZAM_I and 140nt for the *flamenco* region at X:21,769,891..21,789,891 to establish the piRNA density profiles. For comparison between samples, all read counts were normalized by the number of piRNA/siRNA reads used as input (genome-mapped piRNAs or siRNAs, genome-unique piRNAs) and represented in RPM [RPM = read-count * 1,000,000 / (total of genome mapping piRNAs)], or RPKM [RPM = read-count * 1,000,000 / (TE Length/1000) / (total of genome mapping piRNAs)] in the case of density profiles. To assess the ping-pong signature, the counts of 1 to 23nt 5’-overlaps were determined, and the percentage of 10nt 5’-overlaps over the total 1-23nt 5’-overlaps was calculated. The Z-score for 10nt 5’overlaps was determined using the percentages of all 1 to 23nt-long 5’overlaps as a background and was considered significant when it was >1.96. Scatter plots were done with RStudio with antisense piRNAs or siRNAs mapping to the respective internal sequence of Tes (allowing 0-3 mismatches for piRNAs and 0-1 mismatch for siRNAs). All small RNA sequencing done for this study is listed in Supplementary Table S6. Small-RNA sequencing data are available in NCBI GEO database (GSE213368).

### Sterility tests

For the sterility tests, RevI-H2i2 homozygous females were compared to *w*^1118^ females, and homozygous *flamΔZAM* females that carry three functional *ZAM* copies from the RevI-H2 X chromosome were compared to *flamΔZAM* females without these functional *ZAM* copies in their genome. Sterility tests were performed at 25° C. Thirty 3-6-day-old virgin females of each genotype were individually mated with three *w*^1118^ males, and eggs were collected for 24 hours. The number of eggs laid by each female was determined. Then, eggs were kept at 25°C for another 24 hours before determining the egg hatching rate. The experiment was done twice for a total of approximatively 60 replicates for each condition.

### Generation of the *flamΔZAM* line using the CRISPR-Cas9 technology

The *ZAM* copy in the *flamenco* piRNA cluster was deleted using a pCFD6 plasmid that expresses two single guide RNAs (sgRNAs) under the Gal4/UAS system. The sgRNAs were designed to target genome-unique sequences located upstream and downstream of the *flamenco ZAM* copy: TTGTAGCGCTCTTCTTCTCT (sgRNA *flamΔZAM_1*) and AGCGCAACCACGTACAGCGA (sgRNA *flamΔZAM_2*). The pCFD6 plasmid harboring these sgRNAs was injected by the Bestgene company into embryos from the BL#9736 stock (Bloomington stock center) for integration of the plasmid into the genomic site 53B2. The obtained flies were crossed with nanos-gal4; UAS-Cas9 flies (BL#54593, Bloomington stock center). Prior to crossing, the X chromosome of each of these lines was replaced by the X chromosome of the Iso1A *Drosophila* line. After the crosses, 500 lines were derived and screened by PCR using primers specific for the *flamenco ZAM* copy. The detailed crossing schemes are available upon request. Lines without amplification of internal *flamenco ZAM* sequences were selected for PCR amplification of the target locus with primers that framed the *flamenco ZAM* copy and the target sequences of the sgRNAs. The obtained amplicons were sequenced.

## ACKNOWLEDGEMENTS

We thank the *Bloomington Stock Center* for providing stocks. We thank J. Brennecke for providing antibodies. We thank www.flybase.org, “http://www.girinst.org” and NCBI for providing databases and tools. We thank S. Chambeyron and K. Senti for helpful discussions, A. Molaro and the members of the Brasset laboratory for comments on the manuscript. We thank C. Vaury for having established the original RevI-H2 line over 30 years ago, and for preserving it all these years. We thank Y. Renaud and P. Pouchin for helpful advices and development of the iGReD bioinfomatics platform. We also thank the CLIC facility (Clermont Imagerie Confocale). We thank Gentyane Facility for the Quality Control of gDNA.

## AUTHOR CONTRIBUTIONS

Conceptualization, MY, SJ and EB; Validation and Investigation, MY, SMM, AA, NG, BB, Data curation and bioinformatic analyses, MY, SJ, writing—original draft, MY and EB; writing—review and editing, MY, SJ and EB; Supervision, SJ and EB, funding acquisition, MY and EB.

---

**Supplementary Figure 1 (related to Figure 1).**
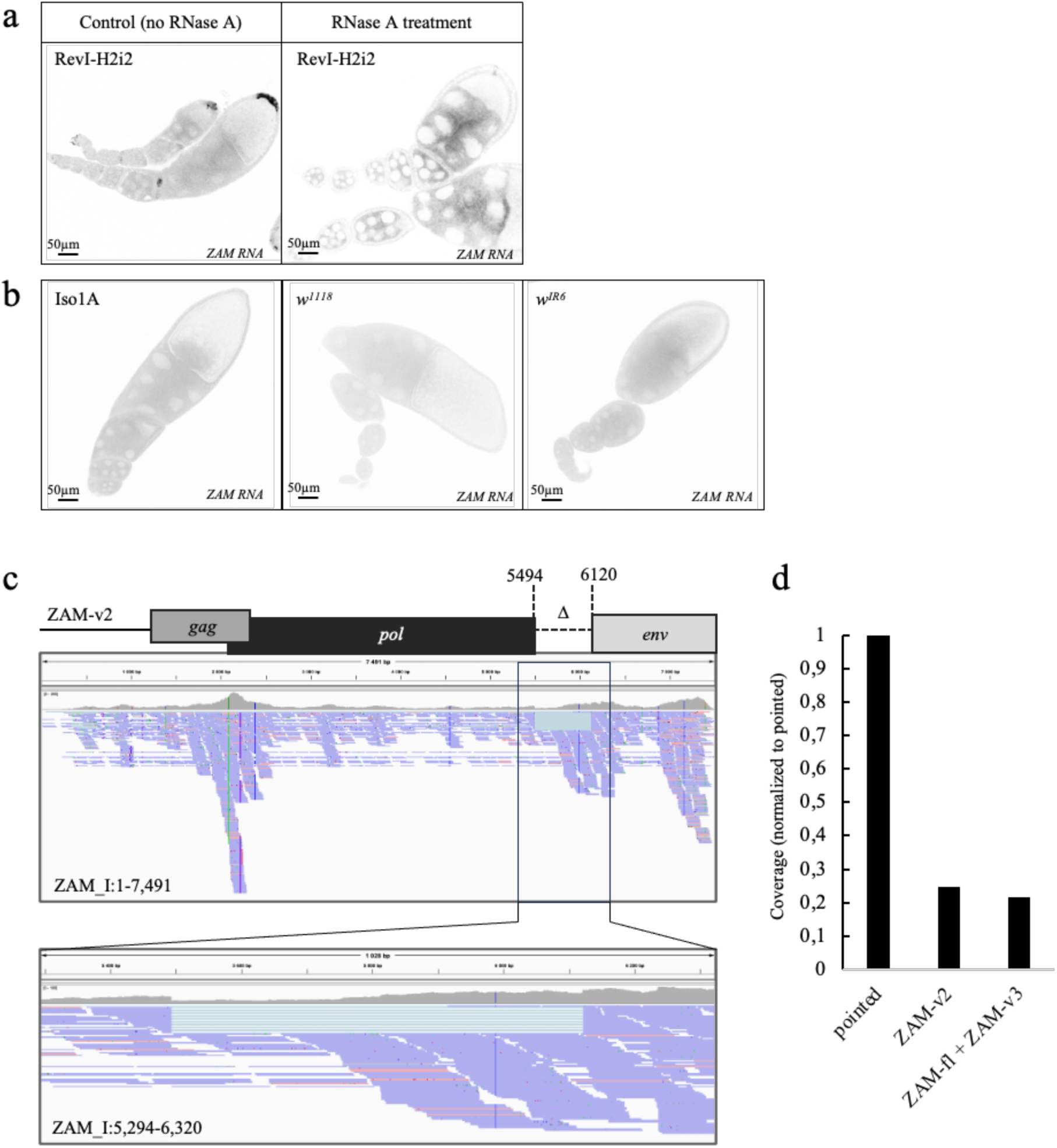
**a.** Color-inverted confocal images of ovarioles from RevI-H2i2 ovaries showing *ZAM* RNA signal without (left panel) and after (right panel) RNase A treatment. **b.** Color-inverted confocal images of ovarioles showing that *ZAM* RNA is completely absent in Iso1A, *w*^1118^, and *w^IR^*^6^ ovaries. **c.** Mapping of mRNA-Seq reads of the full-length *ZAM* (*ZAM-fl*) and of *ZAM-v2* to the reference *ZAM* internal sequences *ZAM*_I (Repbase) allowing splitting of the reads. Above is shown *ZAM-v2* structure. The inset below shows a zoom on the *ZAM-v2* deletion with the corresponding split reads. Reads corresponding to *ZAM* mRNA are in blue, antisense reads are in red. Positions of the shown regions within *ZAM*_I are indicated. d. Coverage of mRNA-seq reads corresponding to the indicated *ZAM*-variants normalized to the average coverage of the gene *pointed*.

**Supplementary Figure 2 (related to Figure 2).**
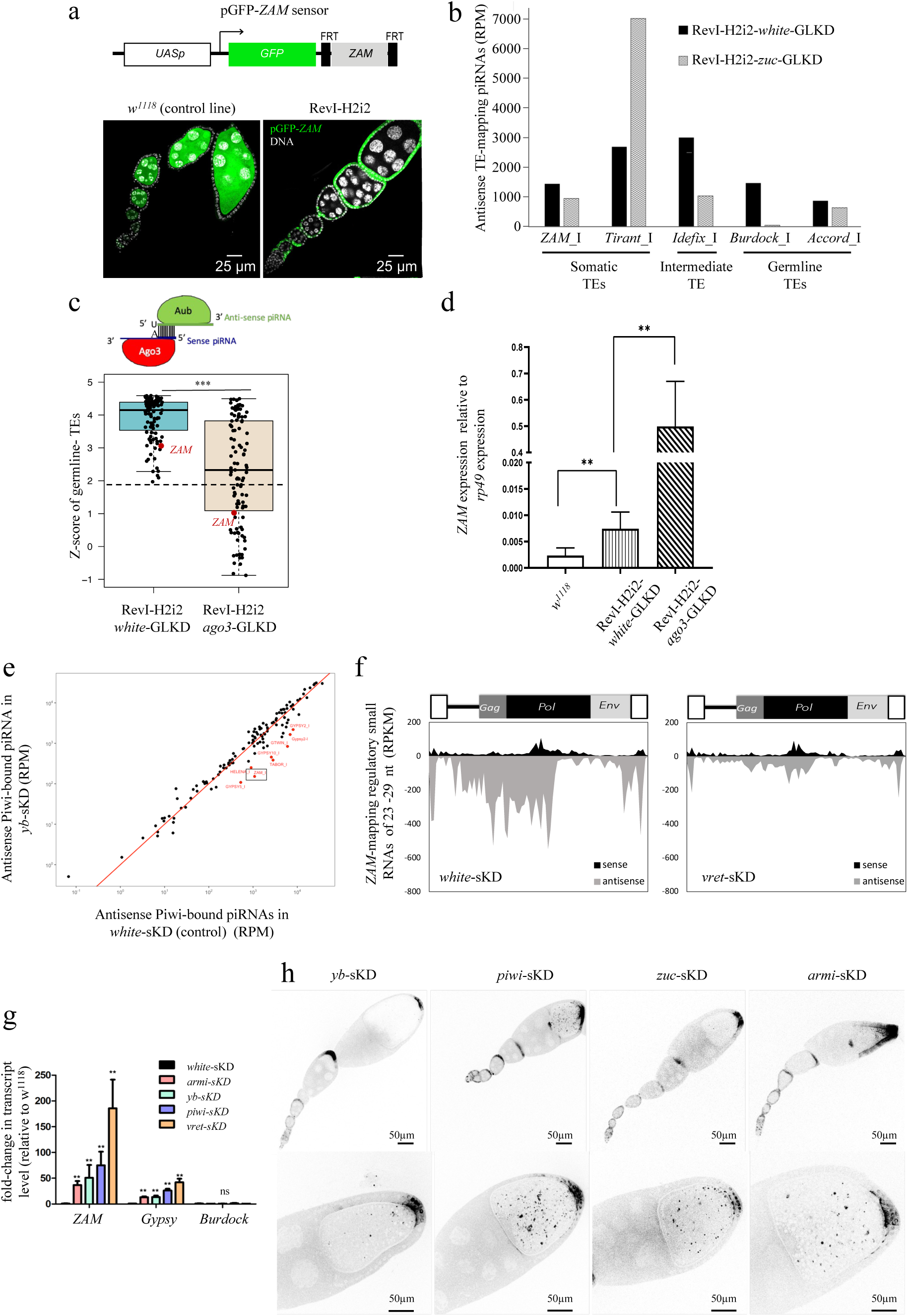
**a**. Confocal images of ovarioles with GFP (green) and DNA (white) staining. Ovarioles of the progeny of a cross between either *w*^1118^ control females or RevI-H2i2 females and males carrying the pGFP-*ZAM* sensor transgene driven by actin-Gal4. **b.** Antisense regulatory piRNAs (23-29 small-RNAs complexed with Argonaute proteins) mapping to TE internal sequences (0-3 mismatches) in the RevI-H2i2 line upon *white-* (control), or *zuc*-GLKD. Normalized per million of genome-mapping piRNAs. **c.** Box plot displaying the Z-score distribution of 10nt 5’-overlaps for TE-mapping piRNAs in the *white*-GLKD and *ago3*-GLKD RevI-H2i2 lines. Each dot represents a TE with a Z-score >1.96 (dotted line) in the control condition (*white*-GLKD). *ZAM* is highlighted in red. The midline indicates the median value, and the box shows the first and third quartile. Error bars indicate the SD. \*\*\**p-value* <0.001 (Mann-Whitney test). **d**. Bar plot showing the steady-state *ZAM* RNA levels in total RNA from ovaries with the indicated genotypes (RT-qPCR, normalized to *rp49*, primer sequences in Supplementary Table S5). At least three biological replicates and two technical replicates were used. \*\**p-value* <0.01 (Mann-Whitney tst); error bars indicate the SD. **e**. Scatter plot showing the normalized counts of antisense Piwi-bound piRNAs mapping to individual internal TE sequence in control ovaries (*white*-sKD) versus *yb*-sKD ovaries. Antisense piRNA counts, mapped allowing up to 3 mismatches, were normalized per million of genome-mapping piRNAs (RPM, here in logarithmic scale). TEs in blue have a *yb*-sKD/*white*-sKD ratio >3; TEs in red have a ratio <0.3. *ZAM* is boxed in black. **f.** Density plot along the *ZAM* sequence of *ZAM*-mapping regulatory piRNAs produced in *white-* and *vret*-sKD ovaries (up to 3 mismatches). **g**. Fold-change in the steady-state *ZAM, Gypsy* and *Burdock* RNA level for the indicated sKD ovaries, compared with *white*-sKD ovaries, measured by RT-qPCR (primer sequences in Supplementary Table S5). At least three biological replicates and two technical replicates were used. \*\**p-value* <0.01 (Mann-Whitney test); error bars indicate the SD. **h**. Color-inverted confocal images of ovarioles (upper panels) and stage 10 egg chambers (lower panels) from the indicated sKD lines showing *ZAM* RNA expression in follicle cells and in the ooplasm in indicated conditions.

**Supplementary Figure 3 (related to Figure 3):**
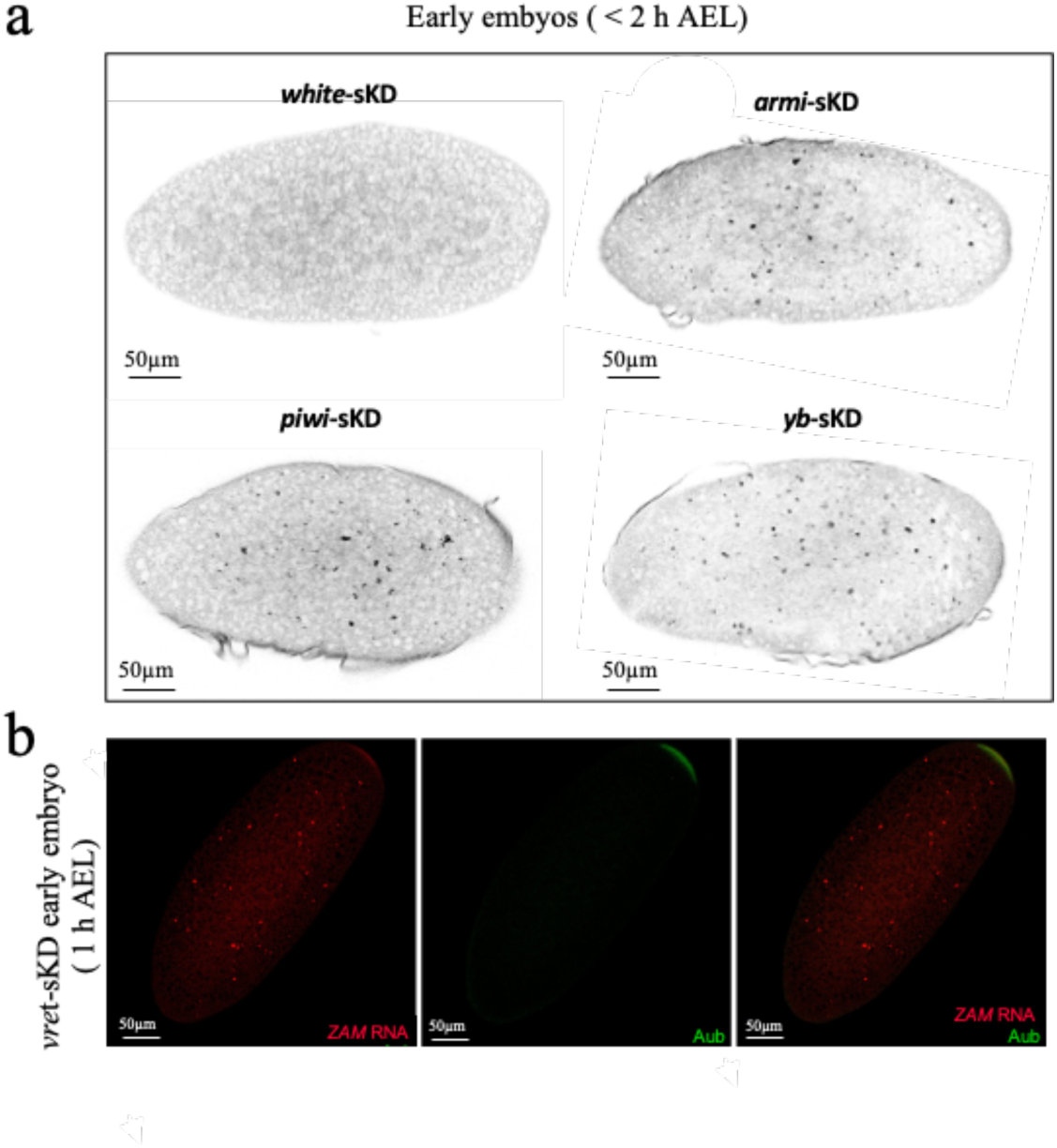
**a.** Color-inverted confocal images of early embryos collected 0-2 hours after egg laying (AEL). Mothers harbor the indicated sKD of the piRNA pathway. *ZAM* RNA signal was detected by smRNA FISH. **b.** Projection of confocal images of stage 1 embryos (0-1 h AEL) laid by *vret*- sKD females showing *ZAM* RNA detected by smRNA FISH and Aub protein immunostaining. Both accumulated at the posterior pole of the embryo where future germ cells cellularize (Z-projection of 25 stacks).

**Supplementary Figure 4.**
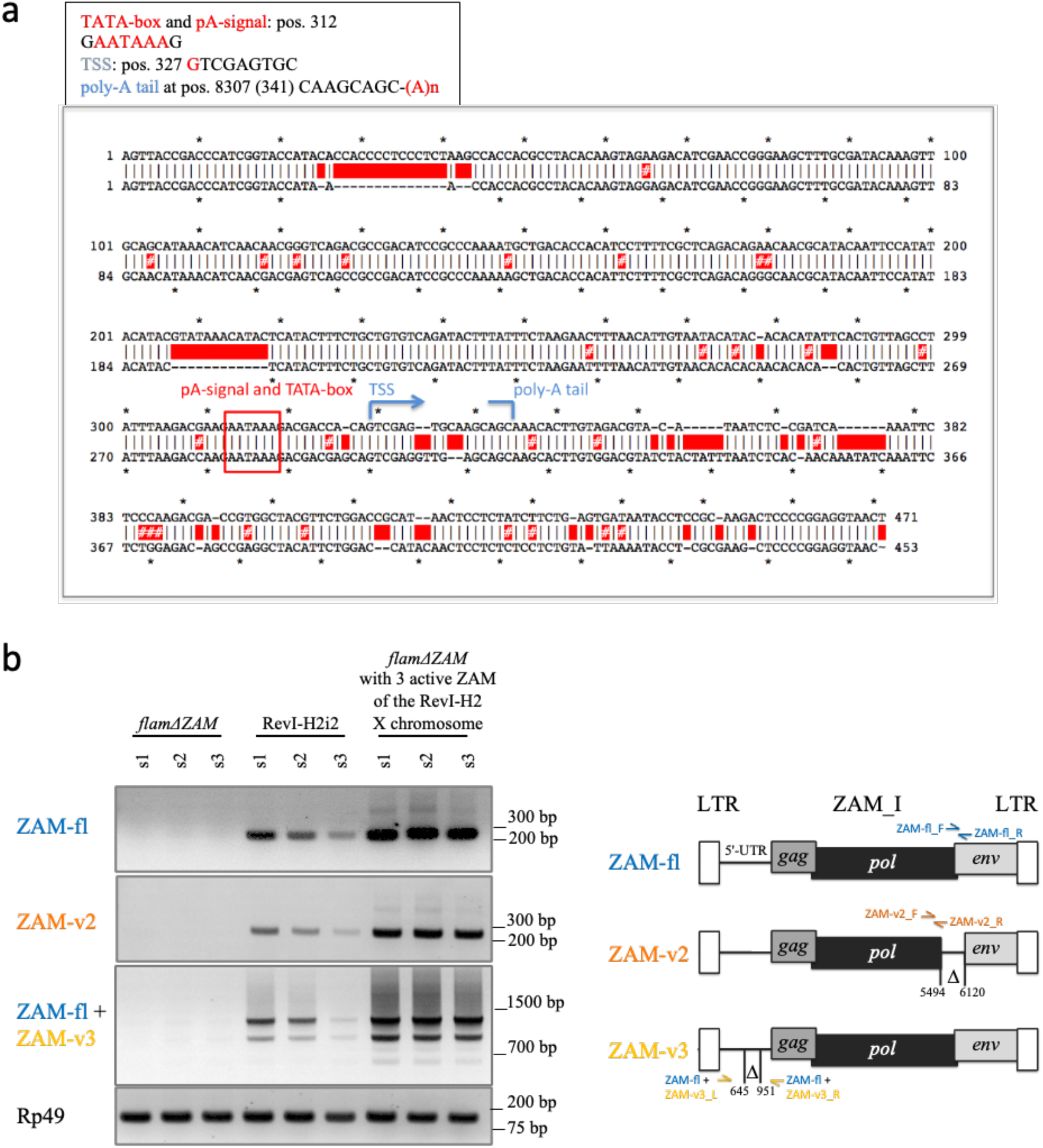
(related to Figure 4): **a.** Sequence alignment of the RevI-H2 consensus *ZAM* LTR and the *ZAM* solo-LTR that remains in the *flamΔZAM* line after CRISPR-Cas9 deletion of the internal *ZAM* sequence in *flamenco*. The image shows the RevI-H2 consensus *ZAM* LTR (upper sequence) and the *ZAM* solo-LTR (lower sequence) with their TATA-box and polyadenylation signal (pA-signal) at position 312 of the *ZAM* consensus, the transcription start site (TSS, at position 327), the start of the poly-A tail at position 341 (CAAGCAGC-(A)n), according to Leblanc et al, 1997. **b.** RT- PCR results showing the expression of the different *ZAM*-variants in the ovaries of the following fly lines: *flam*!i*ZAM*, RevI-H2i2 and *flam*!iZAM with three copies of *ZAM* from the RevI-H2 X chromosome (one *ZAM-fl*, one *ZAM-v2* and one *ZAM-v3*). Three biological replicates were used for each condition (s1, s2, s3). *Rp49* has been used as the housekeeping gene control. The primers used for specific amplification of each variant are indicated on the schematic representation of *ZAM* variants and represented by arrows, and color-coded as follows: Blue: Primers for specific amplification of *ZAM-fl*. Orange: Primers for specific amplification of *ZAM*-v2 (primer *ZAM*-v2_R overlaps left and right edges of the *ZAM*-v2 deletion). Yellow: Primers for amplification of Z*AM-fl* and *ZAM*-v3, where the size of the amplification product differs due to a specific deletion in the 5’-UTR of *ZAM*-v3. (primer sequences in Supplementary Table S5).

**Supplementary Figure 5.**
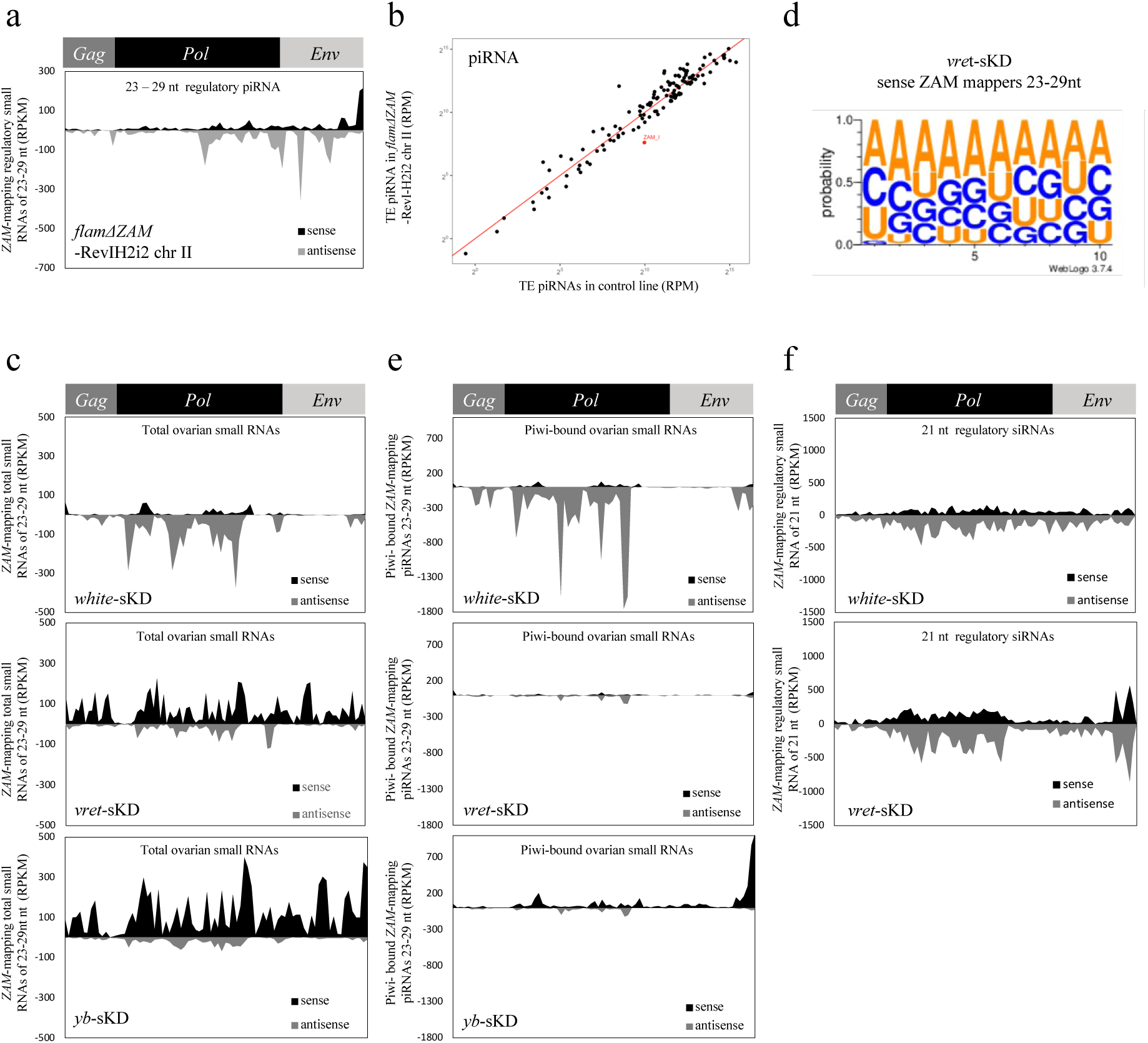
(Related to Figure 5) **a**. Density plot of *ZAM*-mapping regulatory piRNAs (23-29 nt small-RNAs complexed with Argonaute proteins) along the *ZAM* internal sequence in *flamΔZAM*-RevI-H2i2 chr II ovaries (up to 3 mismatches). **b.** Scatter plots showing the normalized counts of regulatory piRNAs mapping to individual internal TE sequences in control ovaries (*white-*sKD) versus *flamΔZAM*- RevI-H2i2 chr II ovaries. Antisense piRNA counts, mapped allowing up to 3 mismatches, were normalized per million of genome-mapping piRNAs (RPM, here in logarithmic scale). **(c)** Density plot of total small RNAs of 23 to 29 nt along the *ZAM* sequence in *white-*sKD, *vret*- sKD and *yb-*sKD ovaries (up to 3 mismatches). **c.** Density plot of total *ZAM*-mapping small RNAs of 23 to 29 nt along the *ZAM* internal sequence in *white-*sKD, *vret*-sKD and *yb-*sKD ovaries (up to 3 mismatches). **d**. Sequence logo for the first ten positions in ZAM-mapping sense small RNAs of 23 to 29nt (from total ovarian small RNA) in *vret*-sKD ovaries. The nucleotide height represents its relative frequency at that position. **e.** Density plot of *ZAM*- mapping Piwi-bound piRNAs along the *ZAM* internal sequence in *white-*sKD, *vret*-sKD and *yb-*sKD ovaries (up to 3 mismatches). **f.** Density plot of *ZAM*-mapping regulatory siRNAs (21nt small-RNAs complexed with Argonaute proteins) along the *ZAM* internal sequence produced in *white*-sKD and *vret*-sKD ovaries (0-1 mismatch).

**Supplementary Figure 6.**
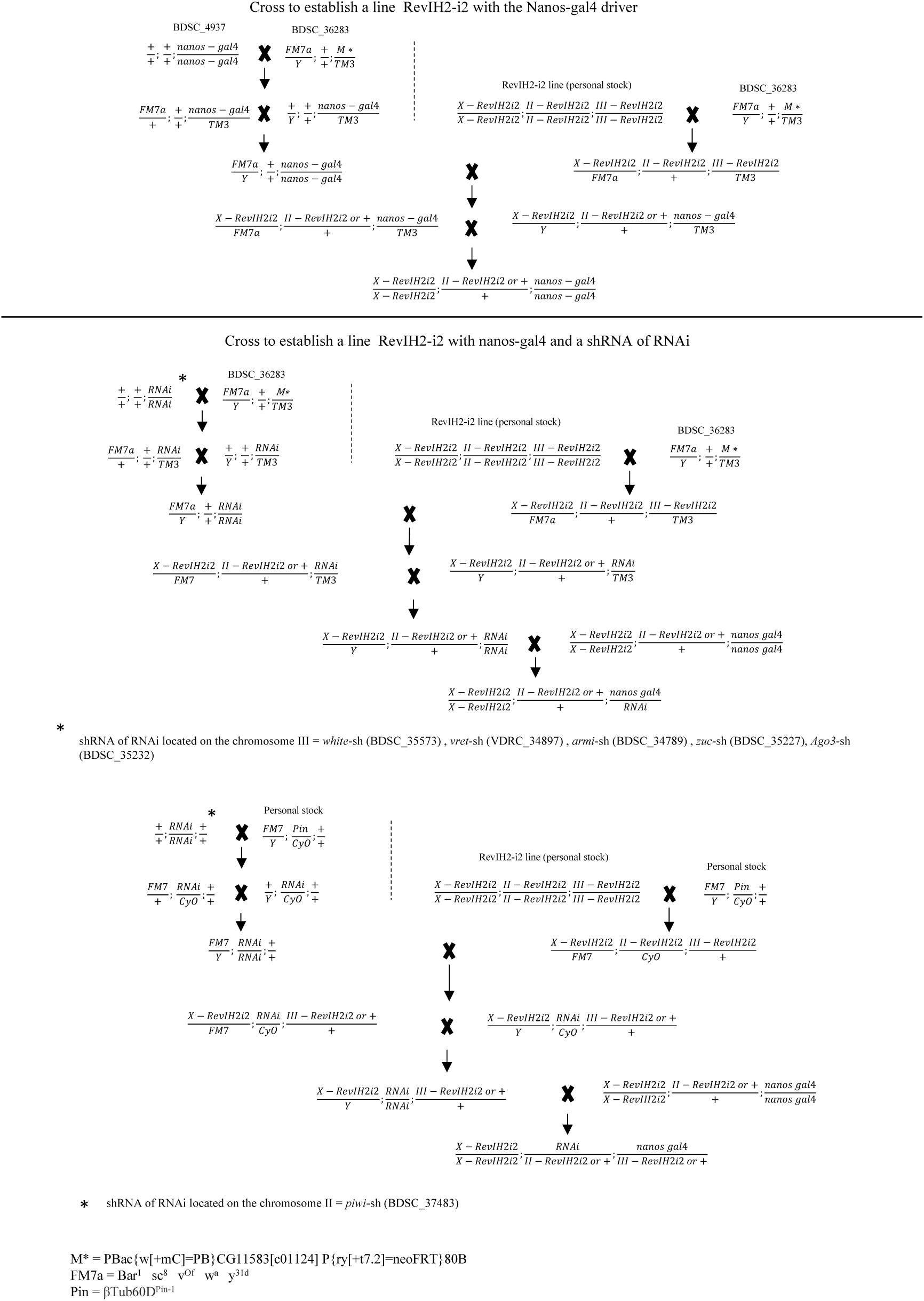
Description of the genetic crosses performed to obtain the different RevI-H2i2-GLKD lines.

**Supplementary Video 1**

Video of confocal Z-stacks overlay of stage 10 egg chambers from vret-sKD ovaries expressing Yolk-protein-1-GFP (green) showing ZAM smRNA FISH signal (red).

**Supplementary Table S1.**
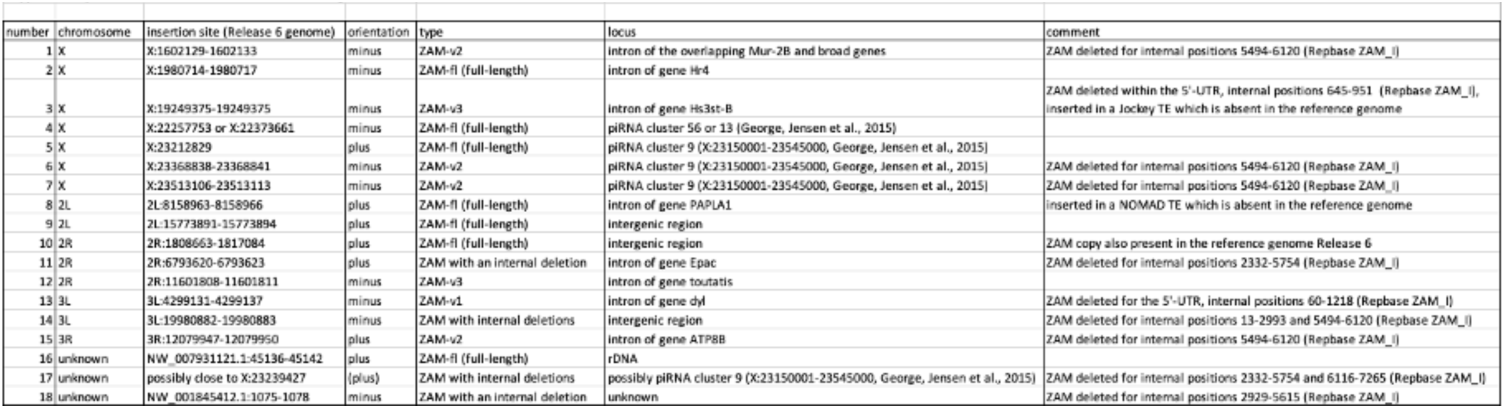
ZAM insertions in the RevI-H2i2 genome

**Supplementary Table 2.**
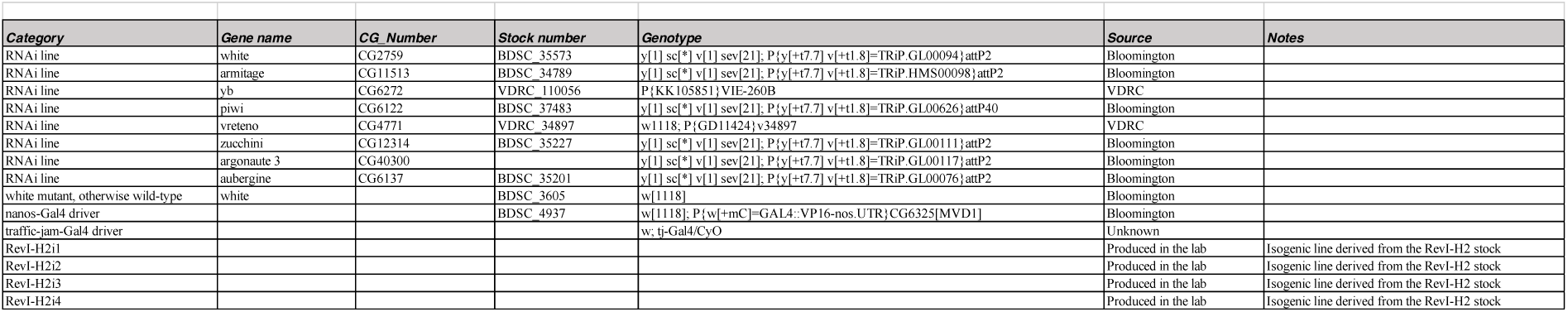
Drosophila lines used

**Supplementary Table 3.**
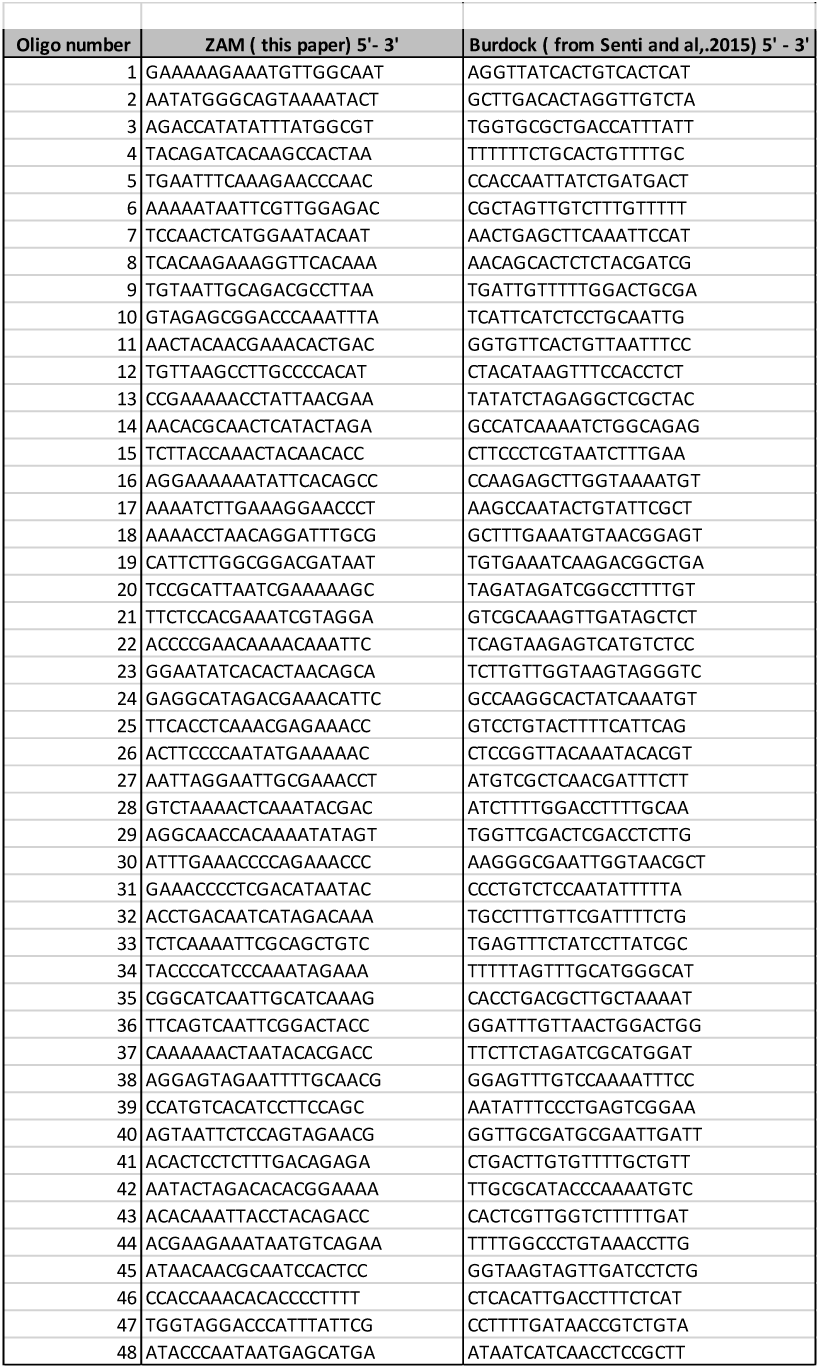
smRNA FISH probes

**Supplementary Table 4.** Antibodies used

| Supplementary Table 4 – Antibodies used |  |  |  |  |  |
| --- | --- | --- | --- | --- | --- |
| Antibody | Source | Dilution | Fluorochrome | Species | Reference |
| Primary antibody |  |  |  |  |  |
| Goat anti-GFP | abcam | 1/1000 |  | Goat | ab5450 |
| Rabbit anti-Aubergine | Homemade (Eurogentec) - polyclonal antibody | 1/1000 |  | Rabbit |  |
| Rabbit anti-Gag-ZAM | Homemade | 1/100 |  | Rabbit |  |
| Rabbit anti Env-ZAM | Homemade | 1/1000 |  | Rabit |  |
| Secondary antibody |  |  |  |  |  |
| Alexa Fluor 488 donkey anti-goat IgG (H+L) | Invitrogen (Fisher) | 1/1000 | Alexa 488 | goat | A11055 |
| Donkey anti-rabbit Alexa 488 IgG (H+L) 2mg/ml | life tech | 1/1000 | Alexa 488 | rabbit | A21206 |
| Goat anti-rat Cy3 | Jackson ImmunoResearch | 1/1000 | Cy3 | rat | 112-165-143 |
| Fluorescent dyes |  |  |  |  |  |
| DAPI | Sigma | 1/500 |  |  | D8417-1MG |

**Supplementary Table S5.**
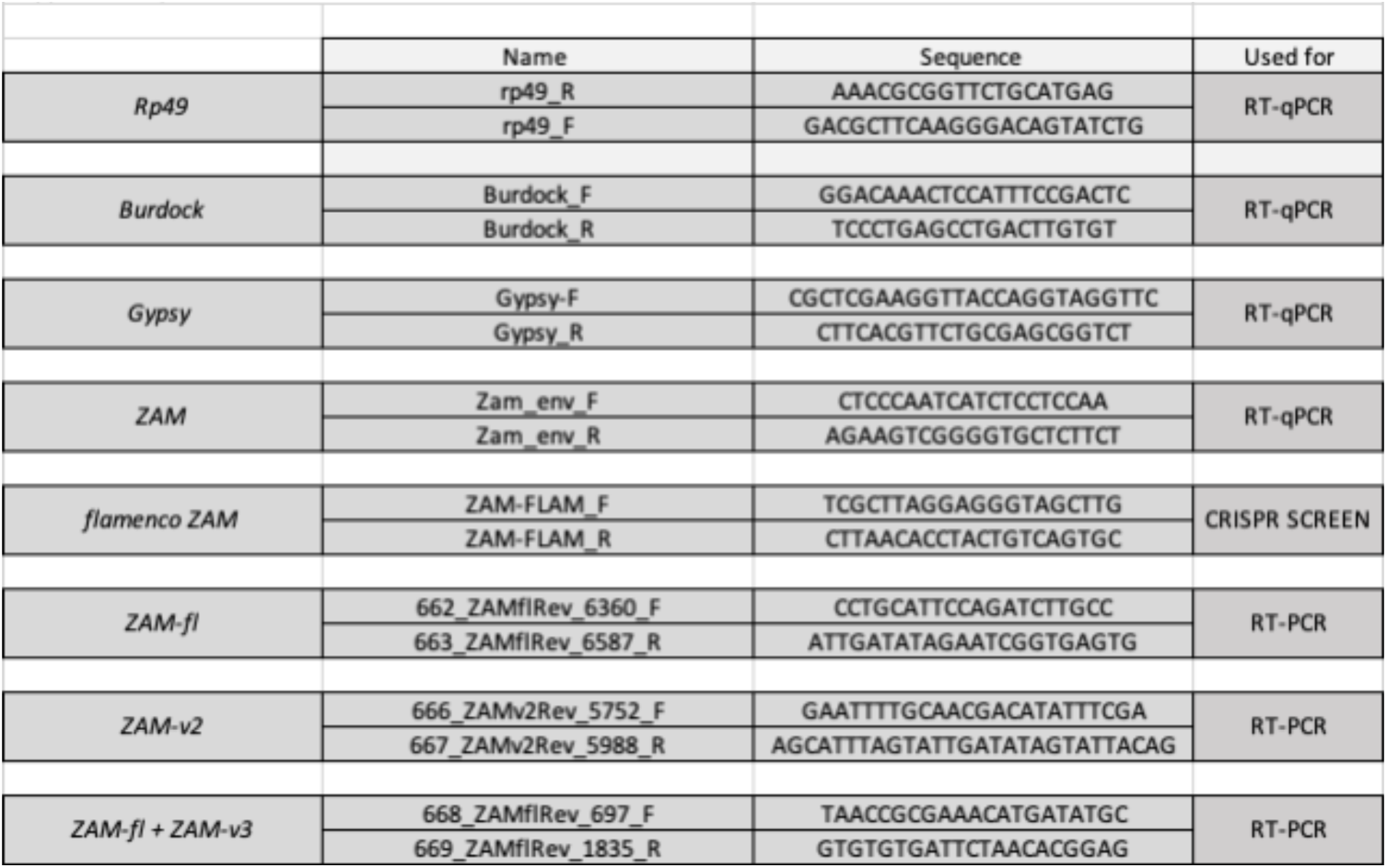
Primers used

**Supplementary Table S6.** small-RNA seq

| Supplementary Table S6 – small-RNA seq |
| --- |
| Small RNA seq |
| TrapR sRNA from Revl-H2i1 |
| TrapR sRNA from Revl-H2i2 |
| TrapR sRNA from Revl-H2i3 |
| TrapR sRNA from Revl-H2i4 |
| Total RNA from Revl-H2i2- nos-Gal4- driven dsRNA-white |
| Total RNA from Revl-H2i2- nos-Gal4- driven dsRNA-Ago3 |
| Total RNA from Revl-H2i2- nos-Gal4- driven dsRNA-Piwi |
| TrapR sRNA- from Revl-H2i2- nos-Gal4- driven dsRNA-Zuc |
| TrapR sRNA- from Revl-H2i2- nos-Gal4- driven dsRNA-white |
| TrapR sRNA from tj-Gal4-driven dsRNA-Vret |
| TrapR sRNA from tj-Gal4-driven dsRNA-white |
| TrapR sRNA from CRISPR flamΔZAM |
| Total RNA from tj-Gal4-driven dsRNA-Vret |
| Total RNA from tj-Gal4-driven dsRNA-white |
| Total RNA from tj-Gal4-driven dsRNA-yb |

## REFERENCES

1. Dewannieux, M. et al. The mouse IAPE endogenous retrovirus can infect cells through any of the five GPI-anchored Ephrin A proteins. PLoS Pathog. 7, e1002309 (2011).

2. Frank, J. A. et al. Evolution and antiviral activity of a human protein of retroviral origin. Science 378, 422–428 (2022).

3. Grow, E. J. et al. Intrinsic retroviral reactivation in human preimplantation embryos and pluripotent cells. Nature 522, 221–225 (2015).

4. Svoboda, P. et al. RNAi and expression of retrotransposons MuERV-L and IAP in preimplantation mouse embryos. Dev. Biol. 269, 276–285 (2004).

5. Tokuyama, M. et al. ERVmap analysis reveals genome-wide transcription of human endogenous retroviruses. Proc. Natl. Acad. Sci. U. S. A. 115, 12565–12572 (2018).

6. Gerdes, P., Richardson, S. R., Mager, D. L. & Faulkner, G. J. Transposable elements in the mammalian embryo: pioneers surviving through stealth and service. Genome Biol. 17, 100 (2016).

7. Nicolau, M., Picault, N. & Moissiard, G. The Evolutionary Volte-Face of Transposable Elements: From Harmful Jumping Genes to Major Drivers of Genetic Innovation. Cells 10, (2021).

8. Ozata, D. M., Gainetdinov, I., Zoch, A., O’Carroll, D. & Zamore, P. D. PIWI-interacting RNAs: small RNAs with big functions. Nat. Rev. Genet. 20, 89–108 (2019).

9. Czech, B. et al. piRNA-Guided Genome Defense: From Biogenesis to Silencing. Annu. Rev. Genet. 52, 131–157 (2018).

10. King, R. C., Aggarwal, S. K. & Aggarwal, U. The development of the female Drosophila reproductive system. J. Morphol. 124, 143–166 (1968).

11. Brennecke, J. et al. Discrete small RNA-generating loci as master regulators of transposon activity in Drosophila. Cell 128, 1089–1103 (2007).

12. Pélisson, A. et al. Gypsy transposition correlates with the production of a retroviral envelope-like protein under the tissue-specific control of the Drosophila flamenco gene. EMBO J. 13, 4401–4411 (1994).

13. Desset, S., Meignin, C., Dastugue, B. & Vaury, C. COM, a heterochromatic locus governing the control of independent endogenous retroviruses from Drosophila melanogaster. Genetics 164, 501–509 (2003).

14. Zanni, V. et al. Distribution, evolution, and diversity of retrotransposons at the flamenco locus reflect the regulatory properties of piRNA clusters. Proc. Natl. Acad. Sci. U. S. A. 110, 19842–19847 (2013).

15. Desset, S., Buchon, N., Meignin, C., Coiffet, M. & Vaury, C. In Drosophila melanogaster the COM locus directs the somatic silencing of two retrotransposons through both Piwi- dependent and -independent pathways. PLoS One 3, (2008).

16. Lau, N. C. et al. Abundant primary piRNAs, endo-siRNAs, and microRNAs in a Drosophila ovary cell line. Genome Res. 19, 1776–1785 (2009).

17. Leblanc, P. et al. Life Cycle of an Endogenous Retrovirus, ZAM, in Drosophila melanogaster. J. Virol. 74, 10658–10669 (2000).

18. Song, S. U., Kurkulos, M., Boeke, J. D. & Corces, V. G. Infection of the germ line by retroviral particles produced in the follicle cells: A possible mechanism for the mobilization of the gypsy retroelement of Drosophila. Development 124, 2789–2798 (1997).

19. Keegan, R. M., Talbot, L. R., Chang, Y. H., Metzger, M. J. & Dubnau, J. Intercellular viral spread and intracellular transposition of Drosophila gypsy. PLoS Genet. 17, 1–22 (2021).

20. Brasset, E. et al. Viral particles of the endogenous retrovirus ZAM from Drosophila melanogaster use a pre-existing endosome/exosome pathway for transfer to the oocyte. Retrovirology 3, 1–9 (2006).

21. Tcheressiz, S. et al. Expression of the Idefix retrotransposon in early follicle cells in the germarium of Drosophila melanogaster is determined by its LTR sequences and a specific genomic context. Mol. Genet. Genomics 267, 133–141 (2002).

22. Sokolova, O. A. et al. Special vulnerability of somatic niche cells to transposable element activation in Drosophila larval ovaries. Sci. Rep. 10, (2020).

23. Malone, C. D. et al. Specialized piRNA Pathways Act in Germline and Somatic Tissues of the Drosophila Ovary. Cell 137, 522–535 (2009).

24. Chalvet, F. et al. Proviral amplification of the Gypsy endogenous retrovirus of Drosophila melanogaster involves env-independent invasion of the female germline. EMBO J. 18, 2659–2669 (1999).

25. Bline, A. P., Le Goff, A. & Allard, P. What Is Lost in the Weismann Barrier? Journal of developmental biology 8, (2020).

26. Leblanc, P., Desset, S., Dastugue, B. & Vaury, C. Invertebrate retroviruses: ZAM a new candidate in D. melanogaster. EMBO J. 16, 7521–7531 (1997).

27. Desset, S. et al. Mobilization of two retroelements, ZAM and Idefix, in a novel unstable line of Drosophila melanogaster. Mol. Biol. Evol. 16, 54–66 (1999).

28. Duc, C. et al. Trapping a somatic endogenous retrovirus into a germline piRNA cluster immunizes the germline against further invasion. Genome Biol. 20, 1–14 (2019).

29. Jurka, J. et al. Repbase Update, a database of eukaryotic repetitive elements. Cytogenet. Genome Res. 110, 462–467 (2005).

30. George, P. et al. Increased production of piRNAs from euchromatic clusters and genes in Anopheles gambiae compared with Drosophila melanogaster. Epigenetics and Chromatin 8, (2015).

31. Czech, B. & Hannon, G. J. One Loop to Rule Them All: The Ping-Pong Cycle and piRNA- Guided Silencing. Trends Biochem. Sci. 41, 324–337 (2016).

32. Meignin, C., Dastugue, B. & Vaury, C. Intercellular communication between germ line and somatic line is utilized to control the transcription of ZAM, an endogenous retrovirus from Drosophila melanogaster. Nucleic Acids Res. 32, 3799–3806 (2004).

33. Atallah, J. & Lott, S. E. Evolution of maternal and zygotic mRNA complements in the early Drosophila embryo. PLoS Genet. 14, e1007838 (2018).

34. Aravin, A. A. & Hannon, G. J. Small RNA silencing pathways in germ and stem cells. Cold Spring Harb. Symp. Quant. Biol. 73, 283–290 (2008).

35. Khurana, J. S. & Theurkauf, W. piRNAs, transposon silencing, and Drosophila germline development. J. Cell Biol. 191, 905–913 (2010).

36. Saito, K. & Siomi, M. C. Small RNA-mediated quiescence of transposable elements in animals. Dev. Cell 19, 687–697 (2010).

37. Klattenhoff, C. et al. Drosophila rasiRNA pathway mutations disrupt embryonic axis specification through activation of an ATR/Chk2 DNA damage response. Dev. Cell 12, 45–55 (2007).

38. Parameswaran, P. et al. Six RNA viruses and forty-one hosts: viral small RNAs and modulation of small RNA repertoires in vertebrate and invertebrate systems. PLoS Pathog. 6, e1000764 (2010).

39. Yu, T. et al. The piRNA Response to Retroviral Invasion of the Koala Genome. Cell 179, 632–643.e12 (2019).

40. Le Rouzic, A. & Capy, P. The first steps of transposable elements invasion: Parasitic strategy vs. genetic drift. Genetics 169, 1033–1043 (2005).

41. Kofler, R., Senti, K. A., Nolte, V., Tobler, R. & Schlötterer, C. Molecular dissection of a natural transposable element invasion. Genome Res. 28, 824–835 (2018).

42. Robillard, É., Le Rouzic, A., Zhang, Z., Capy, P. & Hua-Van, A. Experimental evolution reveals hyperparasitic interactions among transposable elements. Proc. Natl. Acad. Sci. U. S. A. 113, 14763–14768 (2016).

43. Schaack, S., Gilbert, C. & Feschotte, C. Promiscuous DNA: Horizontal transfer of transposable elements and why it matters for eukaryotic evolution. Trends Ecol. Evol. 25, 537–546 (2010).

44. Wierzbicki, F., Kofler, R. & Signor, S. Evolutionary dynamics of piRNA clusters in Drosophila. Mol. Ecol. 1–17 (2021).

45. Yoth, M., Jensen, S. & Brasset, E. The Intricate Evolutionary Balance between Transposable Elements and Their Host: Who Will Kick at Goal and Convert the Next Try? Biology 11, (2022).

46. Mohn, F., Sienski, G., Handler, D. & Brennecke, J. The Rhino-Deadlock-Cutoff complex licenses noncanonical transcription of dual-strand piRNA clusters in Drosophila. Cell 157, 1364–1379 (2014).

47. Luo, S. et al. The evolutionary arms race between transposable elements and piRNAs in Drosophila melanogaster. BMC Evol. Biol. 20, 1–18 (2020).

48. Gebert, D. et al. Large Drosophila germline piRNA clusters are evolutionarily labile and dispensable for transposon regulation. Mol. Cell 1–14 (2021).

49. Shpiz, S., Ryazansky, S., Olovnikov, I., Abramov, Y. & Kalmykova, A. Euchromatic Transposon Insertions Trigger Production of Novel Pi- and Endo-siRNAs at the Target Sites in the Drosophila Germline. PLoS Genet. 10, (2014).

50. Kofler, R. Dynamics of transposable element invasions with piRNA clusters. Mol. Biol. Evol. 36, 1457–1472 (2019).

51. Lerat, E. et al. Population-specific dynamics and selection patterns of transposable element insertions in European natural populations. Mol. Ecol. 28, 1506–1522 (2019).

52. Feschotte, C. & Wessler, S. R. Mariner-like transposases are widespread and diverse in flowering plants. Proc. Natl. Acad. Sci. U. S. A. 99, 280–285 (2002).

53. Navarro-Costa, P. et al. Early programming of the oocyte epigenome temporally controls late prophase i transcription and chromatin remodelling. Nat. Commun. 7, (2016).

54. Bergeron, F., Leduc, R. & Day, R. Subtilase-like pro-protein convertases: from molecular specificity to therapeutic applications. J. Mol. Endocrinol. 24, 1–22 (2000).

55. Klenk, H. D. & Garten, W. Host cell proteases controlling virus pathogenicity. Trends Microbiol. 2, 39–43 (1994).

56. Malik, H. S., Henikoff, S. & Eickbush, T. H. Poised for contagion: evolutionary origins of the infectious abilities of invertebrate retroviruses. Genome Res. 10, 1307–1318 (2000).

57. Hosaka, M. et al. Arg-X-Lys/Arg-Arg motif as a signal for precursor cleavage catalyzed by furin within the constitutive secretory pathway. J. Biol. Chem. 266, 12127–12130 (1991).

58. Guedán, A., Caroe, E. R., Barr, G. C. R. & Bishop, K. N. The Role of Capsid in HIV-1 Nuclear Entry. Viruses 13, (2021).

59. Syomin, B. V., Kandror, K. V., Semakin, A. B., Tsuprun, V. L. & Stepanov, A. S. Presence of the gypsy (MDG4) retrotransposon in extracellular virus-like particles. FEBS Lett. 323, 285–288 (1993).

60. Palmer, K. J. et al. Cryo-electron microscopy structure of yeast Ty retrotransposon virus- like particles. J. Virol. 71, 6863–6868 (1997).

61. Burns, N. R. et al. Symmetry, flexibility and permeability in the structure of yeast retrotransposon virus-like particles. EMBO J. 11, 1155–1164 (1992).

62. Kolliopoulou, A. et al. PIWI pathway against viruses in insects. Wiley Interdiscip. Rev. RNA 10, e1555 (2019).

63. Rozhkov, N. V. et al. Small RNA-based silencing strategies for transposons in the process of invading Drosophila species. RNA 16, 1634–1645 (2010).

64. Rozhkov, N. V. et al. Evolution and dynamics of small RNA response to a retroelement invasion in drosophila. Mol. Biol. Evol. 30, 397–408 (2013).

65. Kawamura, Y. et al. Drosophila endogenous small RNAs bind to Argonaute 2 in somatic cells. Nature 453, 793–797 (2008).

66. Rehwinkel, J. et al. Genome-Wide Analysis of mRNAs Regulated by Drosha and Argonaute Proteins in Drosophila melanogaster. Mol. Cell. Biol. 26, 2965–2975 (2006).

67. Katsuma, S. et al. Transcriptome profiling reveals infection strategy of an insect maculavirus. DNA Res. 25, 277–286 (2018).

68. Petit, M. et al. piRNA pathway is not required for antiviral defense in Drosophila melanogaster. Proceedings of the National Academy of Sciences 113, E4218–E4227 (2016).

69. Quinlan, A. R. & Hall, I. M. BEDTools: a flexible suite of utilities for comparing genomic features. Bioinformatics 26, 841–842 (2010).

70. Grentzinger, T. et al. A universal method for the rapid isolation of all known classes of functional silencing small RNAs. Nucleic Acids Res. 48, (2020).

71. Pogorelcnik, R., Vaury, C., Pouchin, P., Jensen, S. & Brasset, E. sRNAPipe: A Galaxy-based pipeline for bioinformatic in-depth exploration of small RNAseq data. Mob. DNA 9, 4–9 (2018).

